# A ubiquitin-specific, proximity-based labeling approach for the identification of ubiquitin ligase substrates

**DOI:** 10.1101/2023.09.04.556194

**Authors:** Urbi Mukhopadyay, Sophie Levantovsky, Sarah Gharbi, Frank Stein, Christian Behrends, Sagar Bhogaraju

**Affiliations:** European Molecular Biology Laboratory, 71 avenue des Martyrs, 38042, Grenoble, France; Munich Cluster for Systems Neurology, Medical Faculty, Ludwig-Maximilians-Universität München, Munich, Germany; Proteomics Core Facility, European Molecular Biology Laboratory, Heidelberg, Germany

**Keywords:** ubiquitin, E3 ligase, substrate identification, BirA, RING, U box

## Abstract

Ubiquitination of proteins is central to protein homeostasis and other cellular processes including DNA repair, vesicular transport, cell-division etc. The process of ubiquitination is conserved from yeast to humans and is carried out by the sequential action of three enzymes: E1, E2 and E3. There are an estimated >600 E3 ligases in humans that execute ubiquitination of specific target proteins in a spatio-temporal manner to elicit desired signaling effects. Here, we developed a ubiquitin-specific proximity-based labeling method to selectively biotinylate substrates of a given ubiquitin ligase. Our method exploits the proximity and the relative orientation of the E3-ligase catalytic domain with respect to ubiquitin observed in the enzymatic intermediate-state structures of E3-E2∼Ub. By fusing the biotin ligase BirA and an Avi-tag variant to the candidate E3 ligase and ubiquitin, respectively, we were able to specifically enrich *bona fide* substrates and potential new substrates of a ligase using a one-step streptavidin pulldown under denaturing conditions. As proof-of-principle, we applied our method, which we named Ub-POD, to the RING E3 ligase RAD18. RAD18 ubiquitinates DNA-sliding clamp PCNA upon UV-induced DNA damage. We identified PCNA and several other critical players in the DNA damage repair pathway in a single RAD18 Ub-POD experiment. We went on to validate DNA replicase POLE as a possible new substrate of RAD18. Through RAD18 Ub-POD, we were also able to pin down the cellular localization of RAD18-mediated ubiquitination to the damaged DNA nuclear puncta using streptavidin immunofluorescence. Furthermore, we applied Ub-POD to TRAF6, another RING ubiquitin ligase involved in NF-κB signaling and successfully identified known and potentially new TRAF6 substrates. Finally, we adapted our method to the U-box-type E3 ubiquitin ligase CHIP to demonstrate that we can identify substrates of two major classes of mammalian ubiquitin ligases. We anticipate that our method and principle could be widely adapted to all classes of ubiquitin ligases to identify substrates and localize the cellular site(s) of ubiquitination.

## Introduction

Similar to phosphorylation, ubiquitination of proteins has been found to have a role in virtually every cellular process in which it was investigated(*1*). Ubiquitination of substrates is achieved by three enzymes, E1, E2 and E3, acting in a cascade reaction and involves the covalent attachment of the terminal carboxyl group of Ubiquitin (Ub), a 76 amino acid long polypeptide, to substrate lysine through an isopeptide bond. The human genome encodes two E1’s or Ub-activating enzymes, Uba1 and Uba6, that first adenylate the terminal carboxyl group of Ub using an ATP molecule, followed by the formation of a thioester linkage between Ub C-terminus and the catalytic cysteine of E1. There are ∼40 E2’s or Ub-conjugating enzymes which bind to E1 and catalyze a transthioesterification reaction, resulting in an E2∼Ub thioester conjugate. It is estimated that humans encode for >600 E3 ubiquitin ligases which confer specificity and functionality to the Ub cascade by targeted ubiquitination of thousands of cellular substrates(*1*). E3s bind the E2∼Ub conjugate and the substrate to facilitate the transfer of Ub to target lysines of the substrate. The really interesting new gene (RING) family of E3 ligases comprise the majority of E3s in the cell, with an estimated ∼600 members(*2*). RING E3 ligases contain a conserved Zn^2+^-binding RING domain and other auxiliary domains that bind E2∼Ub and substrates, respectively. The RING domain spans ∼70 residues and adopts a conserved alpha/beta topology. In RING E3-catalyzed ubiquitination, Ub is transferred directly from the catalytic cysteine of E2 to the substrate lysine residues. U-box E3 ligases also contain a RING-like catalytic domain that lacks the Zn^2+^-binding site and operate as facilitators of ubiquitination, akin to the RING E3 ligases(*3*). Homologous to E6-AP carboxyl terminus (HECT) family comprises another major class of E3 ligase, with ∼20 members in the human genome. HECT E3 ligases contain a conserved bi-lobal HECT domain at the C-terminus of the protein. The C-lobe of the HECT domain contains an invariant cysteine that accepts Ub from E2∼Ub, before transferring Ub to substrates. RBR (RING-between-RING) type Ub ligases possess two RING domains, one of which contains a catalytic cysteine that becomes transiently thioester linked to Ub, prior to substrate ubiquitination(*4*).

Since E3 ligases are the concluding and decisive actors of all Ub signaling events, substrate identification of a given Ub ligase is critical for gaining insights into its cellular and physiological role. Traditional approaches of identifying protein-protein interactions, such as yeast two-hybrid and immunoprecipitation (IP) coupled proteomics, have provided valuable resources of potential substrates and the cellular pathways a given E3 ligase engages with(*5*). However, in most cases, E3-substrate interactions are transient and difficult to capture by these conventional methods. Many alternate methods have been developed to identify E3 ligase substrates and are reviewed elsewhere (*6*, *7*) in detail. A few such notable approaches are introduced here to provide context for the substrate identification method developed by us. One of the most powerful methods for the identification of E3 ligase substrates uses an antibody that specifically recognizes a diGly remnant that is characteristic of tryptic digested ubiquitinated substrates(*8*). Using quantitative comparison of diGly peptides from cells expressing wildtype (WT) or activity deficient mutant version of E3 ligases, one can identify the substrates and also the target lysines that are modified. Global protein stability (GPS) profiling uses degradation profiles of potential substrates fused to EGFP, coupled with the genetic perturbation of the E3 ligase of interest to identify substrates(*9*, *10*). Another approach, UBAITS uses recombinant fusing of Ub to the E3 ligase of interest to covalently trap the ligase and its cognate substrates, as well as other interacting proteins(*11*). TULIP and TULIP2 methodologies build on the UBAITS principle and use His-tagged Ub-E3 ligase fusion, allowing purification of the E3 ligase-substrate adduct in denaturing conditions(*12*, *13*). E2-dID uses biotin-labeled Ub-E2 conjugates, plus WT or activity deficient mutant E3 ligase of interest, added to the whole cell lysate *in vitro* to identify potential substrates(*14*). An orthogonal ubiquitin transfer (OUT) approach has also been developed using engineered E1, E2 enzymes and E3 ligase of interest that do not cross-react with the endogenous ubiquitin system, thus aiming to specifically target the substrates of the engineered ubiquitin ligase of interest(*15*, *16*). Other notable methods include using a microarray of purified human proteins in an *in vitro* assay(*17*) and a NEDDylator approach, which uses Nedd8-E2 enzyme fused to the E3 ligase of interest(*18*). While no particular method provides a silver bullet solution for the identification of E3 ligase substrates, each method has been proven successful with a specific or a set of ligases and together these methods provide an important toolbox in ubiquitin biology research.

Here we developed an approach to selectively and robustly label the substrates of a given E3 ubiquitin ligase directly in cells. Our approach relies on the conserved but transient intermediate complexes that occur between E2∼Ub and the E3 ligase in the case of RING E3 ligases. Based on the structural information of E2∼Ub-RING E3(*19*, *20*) complexes, we tagged the ligases of our interest with the *E.coli* biotin ligase BirA enzyme and ubiquitin with an acceptor peptide (AP) tag. We co-transfected these constructs in cells, resulting in the proximity and orientation-dependent tagging of Ub (Ub-POD) during the BirA-E3-mediated ubiquitination, leading to biotin-labeled substrates. We first tested this strategy using two unrelated human RING E3 ligases, RAD18 and TRAF6, which are involved in UV-induced translesion DNA synthesis and NF-κB signaling, respectively. We also localized ubiquitination mediated by RAD18 precisely in cells in a Ub-POD experiment using streptavidin immunofluorescence. Next, we successfully employed Ub-POD to a U-box domain containing ligase, CHIP/STUB1 which is involved in the protein quality control machinery by recognizing misfolded proteins and targeting them for degradation. Finally, we delineate the principles and parameters of Ub-POD to facilitate other ubiquitin researchers to adopt Ub-POD and identify the substrates of their favorite ligases.

## Results

### Concept of Ub-POD: a Ub-specific, proximity and orientation-dependent labeling approach for identifying E3 Ub ligase substrates

During Ub transfer to the target proteins, RING E3 ligases bind both substrate and E2∼ubiquitin (E2∼Ub) thioester conjugate, and mediate a direct transfer of ubiquitin from the E2 to the substrate(*21*, *22*). Crystal structures of E2∼Ub-RING E3 complexes have shown that RING E3s first activate the E2∼Ub thioester complex by forming a transient-intermediate complex between E2∼Ub and the RING E3 and subsequently orient the E2∼Ub thioester complex with respect to the substrate(*19*). In the UbcH5A∼Ub-RNF4 complex structure reported by Plechanova *et al.*, 2012, we noted that the N-terminus of the catalytic RING domain and N terminus of Ub point towards the same orientation into solvent and come in close proximity to each other (∼21 Å) (Supplementary Figure 1A). Similarly, analysis of the HECT NEDD4L∼Ub structures(*23*, *24*) has revealed that the C-terminus of the HECT ligase, which harbors the catalytic HECT domain, and the N-terminus of Ub orient towards each other into the solvent (Supplementary Figure 1B). These observations prompted us to develop a precise ubiquitin-specific proximity based labeling approach to identify the substrates of an E3 ubiquitin ligase. Towards this, we tagged the catalytic end of an E3 ubiquitin ligase using the WT *Escherichia coli (E.coli)* biotin ligase enzyme (BirA) and the N-terminus of Ub with BirA’s “acceptor peptide” (AP)(*25*) substrate (Figure 1A). The proximity and the orientation alignment between the E3 catalytic domain and the N-terminus of Ub in the catalytic intermediate complex will enable BirA to catalyze site specific biotinylation of AP linked to Ub. Subsequently, the biotinylated AP-Ub is transferred onto the target proteins, allowing them to be efficiently isolated using streptavidin IP and identified through mass spectrometry (MS) (Figure1 A-B).

**Figure 1:**
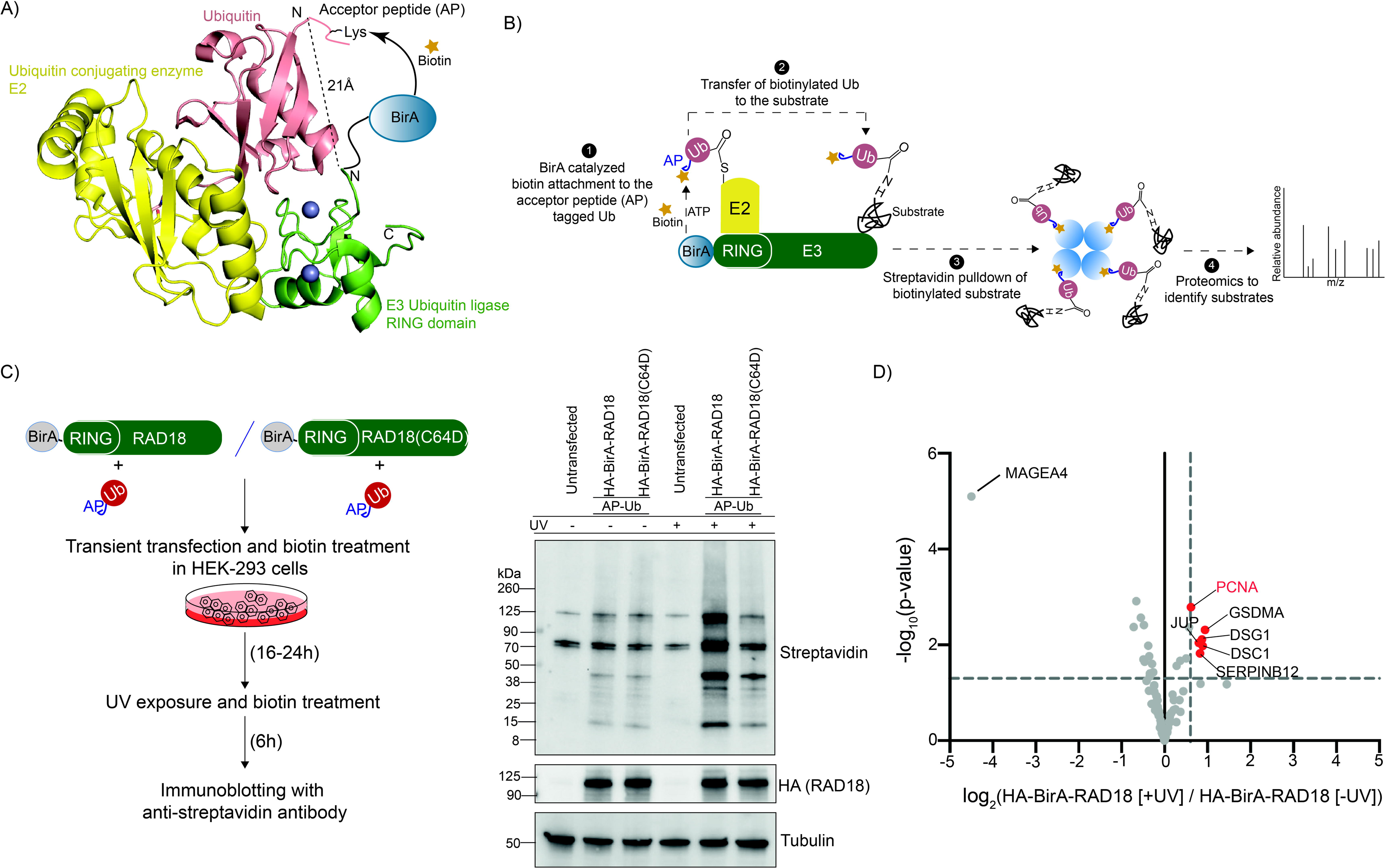
Employing Ub-POD to identify substrates of RING E3 ligase RAD18. A) Structural model of of Ub (salmon) in complex with an Ub conjugating enzyme E2 (yellow) and an Ub ligase E3 (RING) (green). Placing AP-tag at the N-terminus of Ub and BirA at the catalytic end of RING-E3 ligase brings Ap and BirA in proximity (21Å) to each other. B) Schematic representation of the Ub-POD method. C) RAD18 Ub-POD workflow (left panel). HEK-293 cells transfected for 24 h with AP-Ub and HA-BirA-RAD18 WT or HA-BirA-RAD18 C64D were exposed to UV (10 miliJoules/cm2) (right panel). Untreated cells served as control. Lysates were subjected to SDS-PAGE and immunoblotting. D) Volcano plot of proteins labeled by HA-BirA-RAD18 WT in absence or presence of UV exposure (n= 3 biological replicates). Significantly altered proteins (p-value <0.05, log2FC >0.6) are labeled as red circles and known RAD18 substrates are highlighted in red.

### Employing Ub-POD to identify substrates of the RING E3 ligase RAD18

We first tested Ub-POD for substrate identification of the RING E3 ligase RAD18. Upon UV-induced bulky DNA lesions, RAD18, in a complex with the ubiquitin-conjugating enzyme RAD6, mediates monoubiquitination of proliferating cell nuclear antigen (PCNA), thereby triggering translesion synthesis of DNA(*26*). Since RAD18 contains the catalytic RING domain at the N-terminus, we tagged the N-terminus of RAD18 with BirA to generate HA-BirA-RAD18 which we transiently expressed along with AP-Ub in HEK-293 cells in the presence of biotin. After 24 hours, cells were either exposed to UV or left untreated, followed by continued biotin treatment for 6 hours (Figure 1C, left panel, described in detail in the methods section). According to a previous study, maximal ubiquitination of PCNA was only observed 6-8 hours post irradiation (*27*). Cells were collected, lysed and whole cell lysates were subjected to SDS-PAGE followed by immunoblotting with anti-streptavidin antibody. Increased biotinylation clearly occurred in cells treated with UV relative to non-treated cells (Figure 1C, right panel), indicating that Ub-POD can indeed monitor the activity of RAD18. Importantly, introduction of a point mutation (RAD18 C64D) that disrupts the interface of the RAD18-RAD6 complex resulted in substantial reduction in biotinylation. (Figure 1C, right panel). Moreover, expression of BirA alone or BirA-tagged RAD18 without AP-Ub only resulted in negligible biotinylation as compared to co-expression of these proteins together with AP-Ub (Supplementary Figure 1C). Likewise, co-expression of AP-Ub with BirA alone only led to minimal biotinylation (Supplementary Figure 1C). Together these results indicated that BirA on its own does not notably biotinylate AP-Ub compared to catalysis-induced proximity between BirA-RAD18 and AP-Ub. Prompted by the positive outcome, we next employed proteomics to identify these biotinylated proteins, which are possible ubiquitination substrates of RAD18. We followed similar experimental conditions as above and the biotinylated proteins were immunoprecipitated (IPed) using streptavidin beads under denaturing conditions from both UV treated and untreated cells in triplicates (see methods). Quantitative MS analysis of the IPed fraction was performed using TMTplex labeling. We identified PCNA as the most enriched candidate (Figure 1D, Supplementary table 1) in the UV-treated condition, providing proof of principle that the Ub-POD method can be employed to identify substrates of a RING ubiquitin ligase. Intriguingly, Melanoma associated antigen 4 (MAGEA4) was found enriched in the untreated condition (Figure 1D, Supplementary table 1). MAGEA4 has been recently identified as the stabilizer of RAD18 and was also shown to be ubiquitinated by RAD18 *in vitro* (*28*). Our data indicates that RAD18 likely ubiquitinates MAGEA4 when cells are not exposed to UV but preferentially ubiquitinates PCNA post UV-exposure. In the next step, we sought to improve the specificity and sensitivity of substrate identification using Ub-POD to increase the number of identified proteins and to improve the enrichment of substrates.

### A variant of the AP tag improves the sensitivity of Ub-POD

Due to a relatively high intrinsic affinity of AP for BirA, protein-protein interaction (PPI) independent biotinylation may occur in cells, giving rise to background biotinylation of free AP-Ub that is not primed for ubiquitination by the BirA-tagged Ub ligase. Fernandez Suarez *et al.,* 2008 tested multiple variants of the AP sequences for their ability to specifically get biotinylated strictly dependent on the proximity to the BirA labeled protein(*29*). Some of these variants included the deletion of up to three amino acids from one or both ends of AP to decrease its interaction surface area with BirA. Based on this, we prepared two variants of AP-Ub; (-2)AP-Ub in which two N-terminal residues from AP were removed and AP(-3)-Ub in which C-terminal 3 residues of AP were deleted (Figure 2A). WT as well as truncated AP-Ub constructs were overexpressed in HEK-293 cells, along with BirA or BirA-RAD18, followed by biotin treatment and UV exposure (Figure 2A). In agreement with previous reports, both truncated AP-variants, (-2)AP-Ub and AP(-3)-Ub showed much lower background biotinylation, in presence of BirA, compared to AP-Ub. While AP-Ub as well as both variants showed increased biotinylation in presence of BirA-RAD18 compared to BirA, the relative increase in biotinylation between BirA-RAD18 and BirA alone was more prominent in the presence of (-2)AP-Ub or AP(-3)-Ub than with AP-Ub (Figure 2A). This indicated that both (-2)AP-Ub and AP(-3)-Ub are more suitable for Ub-POD compared to AP-Ub. Fernandez Suarez *et al.,* 2008 showed that the proximity-dependent biotinylation of AP(-3) is less pronounced due to increased K_M_ (350µM) of AP(-3) compared to AP (K_M_=25µM), indicating that AP(-3) binds much weaker to BirA. AP(-3) also exhibited a diminished k_cat_ (0.5 min^-1^) compared to AP (14 min^-1^) and it was proposed that AP(-3) tag is only suitable to detect protein-protein interactions with a half-life of more than a minute(*29*). Since ubiquitination reactions are typically much faster than the kinetics of AP(-3) biotinylation by BirA(*25*, *29–31*), we chose (-2)AP-Ub for our subsequent Ub-POD experiments. When (-2)AP-Ub was used in Ub-POD of RAD18, the overall number of identified hits increased in the MS analysis compared to the experiment performed with AP-Ub (Figure 2B, Figure 1D, Supplementary table 2). BirA and (-2)AP-Ub co-expression was used as a control in this experiment as it showed negligible background biotinylation (Figure 2A). Gene ontology (GO) enrichment analysis of RAD18 Ub-POD hits highlighted GO terms related to base excision repair (p≤0.001), nucleotide excision repair (p≤0.001), mismatch repair (p≤0.01) and DNA replication (p<0.001) which are highly relevant to the role of RAD18 in UV-induced DNA damage repair (Figure 2C). Since RAD18 C64D showed considerably lower background biotinylation compared to RAD18 WT (Figure 1C), we next probed if RAD18 C64D can serve as a control in RAD18 Ub-POD experiments. Surprisingly, only four proteins were identified as significant hits in RAD18 WT compared to RAD18 C64D (Supplementary Figure 2A). The enrichment of (log2 Fold Change (FC)) PCNA in RAD18 C64D was only slightly lower compared to RAD18 WT. As a result, PCNA was not identified as a significant hit in the comparison between RAD18 WT and RAD18 C64D (Supplementary figure 2A-B, Supplementary table 2) but surprisingly, PCNA was found enriched in Ub-POD experiment performed with RAD18 C64D (Supplementary Figure 2B). Since RAD18 is known to dimerize(*32*), exogenously expressed HA-BirA-RAD18 C64D might dimerize with endogenous RAD18, making this mutant a suboptimal control for proximity-dependent proteomics experiments. Since many RING E3 ligases are known to dimerize(*33*), the use of activity-deficient point mutants as controls in Ub-POD experiments will need to be carefully considered and compared to BirA alone.

**Figure 2:**
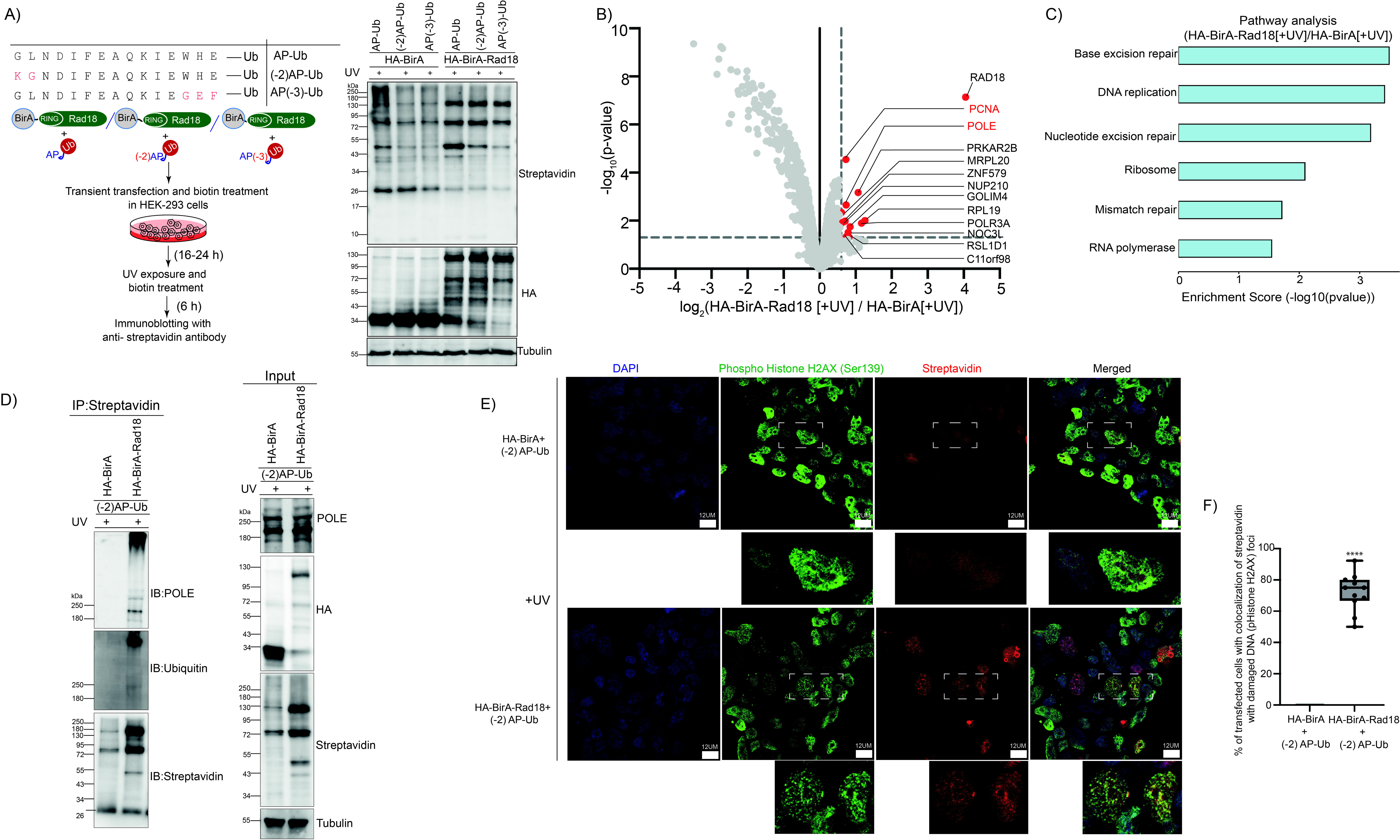
A variant of AP-tag increases the efficiency of Ub-POD. A) Amino acid substitutions of different AP-tag variants compared to AP WT (left, upper panel). Workflow for AP-tag optimization (left, lower panel). AP WT and AP-tag variants were transfected for 24 h in HEK-293 cells along with HA-BirA or HA-BirA-Rad18 followed. Biotin was added at the time of transfection and cells were recovered 6h post UV irradiation. Lysates were subjected to SDS-PAGE and immunoblotting. B) HEK-293 cells transfected with (-2)AP-Ub and HA-BirA or HA-BirA-RAD18 followed by 6 h UV exposure. Cells were kept in 100 µM biotin the whole time. Lysates were subjected to streptavidin IP followed by MS analysis. Volcano plot of proteins labeled by HA-BirA and HA-BirA-RAD18 (n= 3 biological replicates). Significantly altered proteins. Proteins with p-value <0.05, log2FC >0.6 are labeled as red circles. Known and potential RAD18 substrates are highlighted in red. C) GO term analysis of RAD18 substrate candidates. Bar graph shows significantly enriched pathways. D) HEK-293 cells transfected with (-2)AP-Ub and HA-BirA or HA-BirA-RAD18 were treated with biotin and exposed to UV as in B. Lysates were subjected to streptavidin IP. 5 % of lysates were kept as input. IP and input samples were separated by SDS-PAGE followed by immunoblotting with specific antibodies. E) Confocal microscopy of HEK-293 cells transfected with (-2)AP-Ub and HA-BirA or HA-BirA-Rad18 were fixed after indicated treatments and immunostained with anti-phospho-histone H2AX (Ser 139) (green) and anti-streptavidin (red) antibodies. Scale bar, 12 µm. F) Percentage of transfected cells showing colocalization of streptavidin and phospho-histone H2AX signals (****p<0.0001) (n= 10 different fields).

Among the identified RAD18 Ub-POD hits (Figure 2B), DNA polymerase epsilon catalytic subunit (POLE) was of particular interest to us, as this is a predicted substrate of RAD18 with a high confidence score in Ubibrowser(*5*, *34*). POLE is crucial for optimal leading-strand DNA synthesis and POLE-mediated leading-strand replication is dependent on the sliding-clamp processivity factor PCNA(*35*). In the presence of DNA lesions, monoubiquitination of PCNA by RAD18 upon replication-stress results in switching of replicative polymerases (POLE and POLD) with translesion synthesis polymerases. A previous diGly proteomics analysis aiming to identify DNA damage-induced ubiquitination events revealed that ubiquitination of POLE is upregulated upon UV exposure (*36*). To validate that POLE is indeed ubiquitinated by RAD18 upon UV exposure, we overexpressed either BirA or BirA-RAD18 in HEK-293 cells along with (-2)AP-Ub and treated these cells with UV. Immunoblotting of the streptavidin-IPed fraction with antibodies recognizing POLE and Ub showed the appearance of polyubiquitinated POLE species only when BirA-RAD18 is expressed (Figure 2D). Hence, apart from monoubiquitinating PCNA upon UV-induced DNA damage, RAD18 might also be involved in polyubiquitination of POLE. Since RAD18 is known to monoubiquitinate its substrates, it remains to be seen whether RAD18 alone can polyubiquitinate POLE or it does so in conjunction with other E3 ligases. Future studies are required to understand the exact nature and significance of this intriguing new ubiquitination event in the context of the various DNA repair pathways that RAD18 plays a role in(*27*, *37–39*).

### The effect of Ub recycling on Ub-POD

The process of ubiquitination is reversible through the action of the deubiquitinase (DUB) family of enzymes(*40*, *41*). This could potentially lead to the recycling of biotinylated (-2)AP-Ub employed in the Ub-POD experiment, potentially jeopardizing the specificity and efficiency of substrate identification. To understand the impact of Ub recycling on the Ub-POD method, we used a nonselective, broad spectrum DUB inhibitor PR619 (2,6-diaminopyradine-3,5-bis(thiocynate)) that efficiently silences four of the five DUB families(*42*). We treated BirA-Rad18 transfected cells with 10 µM of PR619, 2 hours post UV exposure. The control BirA transfected cells were not treated with PR619. Proteomics analysis showed that PR619 treatment increases the efficiency of PCNA identification compared to non-treated cells (Supplementary figure 3A-B, Supplementary table 3). PR619 treatment also increased the overall number of proteins identified as hits and GO annotation analysis showed that there is enrichment of proteins involved in the process of DNA replication (p≤0.01) (Supplementary Figure 3B-C). We identified several proteins which are predicted to be RAD18 substrates by the repositories such as Ubibrowser(*5*), BioGRID(*43*), Reactome(*44*) and these candidates have been highlighted in supplementary table 3. Our data indicates that a pan DUB inhibitor treatment could improve substrate identification by Ub-POD as executed here for RAD18.

### Exploiting Ub-POD to monitor spatial ubiquitination

Visualizing where in the cell ubiquitination of substrates is taking place adds cellular context to understanding the function of a given ligase. Ub-POD experiments coupled to streptavidin immunofluorescence and confocal microscopy can in principle reveal the cellular localization of ubiquitinated substrates. Taking advantage of the fact that RAD18-mediated PCNA monoubiquitination and the following polymerase switching event take place at UV-irradiation induced RAD18 nuclear foci(*27*), we addressed whether Ub-POD can visualize PCNA ubiquitination at the damaged DNA foci. In irradiated cells expressing BirA-RAD18 and (-2)AP-Ub, streptavidin co-localized with phosphorylated Histone H2AX (Ser139) (γ-H2AX), which is a *bona fide* marker of ionizing radiation induced DNA damage(*45*) (Figure 2E-F). As expected, streptavidin and HA (detects BirA or BirA-RAD18) co-localized in nuclear puncta only in BirA-RAD18 expressing cells (Supplementary Figure 4). Together these results indicated that Ub-POD can indeed provide insights into the cellular localization of a given E3 ligase-mediated ubiquitination reaction.

### Employing Ub-POD on the RING E3 ligase TRAF6

We next sought to benchmark our approach with the well-studied RING E3 ligase TNF receptor-associated factor 6 (TRAF6) which mediates K63 ubiquitin linkage types in concert with the ubiquitin-conjugating enzyme heterodimer UBE2N-UBE2V1. TRAF6 bridges tumor necrosis factor (TNF) receptor activation via exogenous agents or endogenous mediators and subsequent activation of transcriptional responses via NF-κB and MAPK pathways (*46*, *47*). Firstly, we performed a biotinylation time course to shorten biotin incubation and reduce non-specific labeling. We transiently co-transfected HEK-293 cells with (–2)AP-Ub and BirA-TRAF6 or BirA alone for 24 h and grew cells for up to 24 h in the presence of biotin. While BirA was expressed at substantially higher levels than BirA-TRAF6, cells expressing the former did not show any biotinylation irrespectively of the biotin incubation time. In contrast, biotinylation was clearly detectable in BirA-TRAF6 expressing cells 15 min after biotin addition (Figure 3A). Since signaling downstream of TRAF6 can be observed between 5 and 20 min after pathway activation(*48*), we settled on 15 min for all future labeling experiments to capture ubiquitination substrates.

**Figure 3:**
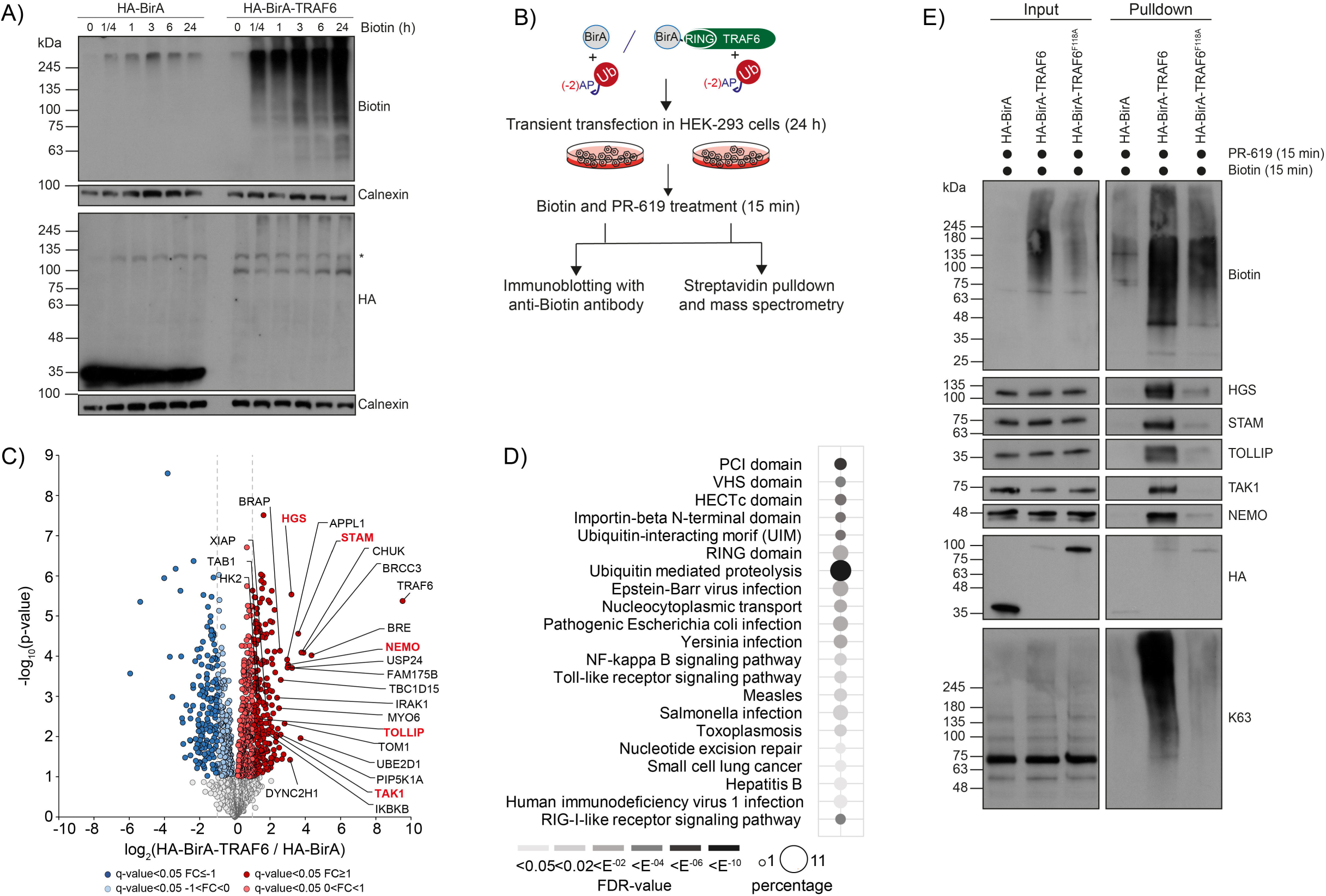
Employing Ub-POD on the RING E3 ligase TRAF6. A) HEK-293 cells were transfected with (-2)AP-Ub and HA-BirA or HA-BirA-TRAF6 for 24 h. Biotin (100 µM) was added for indicated time points. B) Workflow for TRAF6 Ub-POD MS. HEK-293 cells transfected for 24 h with (-2)AP-Ub and HA-BirA or HA-BirA-TRAF6 were incubated with biotin (100 µM) and PR-619 (10 µM) for 15 min prior to cell harvest. Lysates were subjected to streptavidin IP and MS analysis. C) Volcano plot of proteins labeled by HA-BirA-TRAF6 or HA-BirA (n = 4 biological replicates). Significantly altered proteins are shown in dark red or blue (FDR <0.05, log2FC >I1I) and light red or blue (FDR <0.05, 0 > I log2FC I < I1I) (two-sided t-test). Known and potential TRAF6 substrates are labeled in black. Substrates validated in E are highlighted in red. D) GO terms of proteins found to be significantly enriched in HA-BirA-TRAF6 versus HA-BirA with FC ≥1. Dot size correlates to number of proteins; dot color to GO term enrichment (FDR value). E) Validation of TRAF6 substrates identified via TRAF6 Ub-POD MS. HEK-293 cells transfected for 24 h with (-2)AP-Ub and HA-BirA, HA-BirA-TRAF6 WT or HA-BirA-TRAF6 F118A were incubated with biotin (100 µM) and PR-619 (10 µM) for 15 min and subjected to denaturing streptavidin IPs followed by immunoblotting.

To further examine the specificity of the Ub-POD method, we compared biotinylation levels of TRAF6 WT, the dimerization mutant TRAF6 F118A and the catalytically dead mutant TRAF6 C70A(*49*, *50*). As expected, both mutants showed a substantial decrease in biotinylation despite their higher expression compared to TRAF6 WT (Supplementary Figure 5A). We speculate that residual biotinylation might stem from endogenous TRAF6 which might dimerize with overexpressed TRAF6 mutants.

To identify potential TRAF6 substrates, we transiently co-transfected HEK-293 cells with (-2)AP-Ub and N-terminally tagged BirA-TRAF6 or BirA alone for 24 h, and subsequently treated cells with biotin and PR-619 for 15 min (Figure 3B). Following lysis, an aliquot was removed from each replicate to confirm induction of biotinylation (Supplementary Figure 5B). Notably, overexpression of TRAF6 was sufficient to induce downstream NF-κB signaling, possibly due to enhanced dimerization of TRAF6 which is crucial for its autoubiquitination and catalytic activity(*49*). We subjected these samples to streptavidin IP under denaturing condition, on-beads tryptic digestion and MS analysis. Across the different conditions, quadruplicate samples showed high Pearson correlation (r ≥ 0.928). In total, we identified 2,145 proteins of which 1,331 proteins passed stringent filtering. 503 proteins were significantly enriched in BirA-TRAF6 expressing cells compared to HA-BirA control expressing cells. 231 of these proteins reached the threshold of FDR <0.05 and fold change (FC) >1 (Figure 3C, Supplementary table 4, 5). GO annotation analysis of proteins reaching the threshold of FDR<0.05 and FC >1 (231 proteins) unveiled shared protein motifs, such as E3 ligase domains (RING and HECT) and ubiquitin-interacting motifs (UIM). Intriguingly, positively regulated hits showed specific enrichment of proteins involved in immune response pathways against bacterial and viral infection, including Toll-like receptor and NF-κB signaling pathway, amongst others, which gives us confidence in the robustness of our approach even in non-immune cell types such as HEK-293 cells (*47*, *50*) (Figure 3D). TRAF6 is the strongest regulated hit due to its autoubiquitination in the active, oligomerized state(*49*). Furthermore, our approach enabled the specific enrichment of *bona fide* TRAF6 substrates (NEMO, IRAK1, TAK1) (*47*, *50–52*) (*50*, *53*). Interestingly, we also enriched for the deubiquitinating enzyme complex (BRISC), which is specific for lysine 63-linked ubiquitin hydrolysis (*54*). Furthermore, strongly regulated hits are VHS-domain containing and Ub-binding proteins located at endosomes (STAM, HGS, TOM1), which are involved in receptor tyrosine kinase (RTK)-mediated endocytosis (*55*). Next, we sought to validate known and potential new TRAF6 substrates. Candidates were labeled in cells by BirA-TRAF6 and enriched via streptavidin IP prior to their detection by immunoblotting. Notably, BirA-TRAF6 F118A and BirA only expressing cells served as negative controls. Using this approach, we were able to confirm known TRAF6 substrates such as NEMO and TAK1 whose biotinylation was absent in BirA only or drastically reduced in BirA-TRAF6 F118A expressing cells despite substantial higher expression levels of these control proteins (Figure 3E). Importantly, we also confirmed TRAF6 labeling-dependent enrichment of the ESCRT proteins STAM and HGS as well as of the selective autophagy receptor TOLLIP, all of which were identified as potential new substrates of TRAF6 (Figure 3E). In addition, we observed strong enrichment of K63-linked polyubiquitylated proteins, suggesting that substrate candidates might be modified by this TRAF6 signature ubiquitin chain type.

### Employing Ub-POD to identify substrates of the U-box-containing ligase CHIP

Next, we asked if Ub-POD can be applied to a U-box domain containing Ub ligase. The U-box domain adopts the same fold as the RING domain but lacks the Zn^2+^ coordination residues which are replaced by a tight hydrogen bonding network, contributing to the integrity of the catalytic domain structure(*56*, *57*). CHIP (carboxyl terminus of Hsc70-interacting protein) which is also called STUB1 (STIP1 homology and U box-containing protein 1) is a U-box domain containing Ub ligase that functions as a key regulator of the protein quality control machinery. CHIP binds the molecular chaperones Hsc70, Hsp70 and HSP90 through its N-terminal tetratricopeptide repeat (TPR) domain and facilitates the polyubiquitination of misfolded client proteins via its C-terminal catalytic U-box(*58–60*). We employed Ub-POD on CHIP to benchmark our approach for a U-box E3 ligase and to possibly identify new substrates of CHIP. In line with the concept of Ub-POD, we prepared a CHIP construct where BirA is tagged to the C-terminus of CHIP (CHIP-BirA), since the catalytic U-box domain of CHIP is located at the extreme C-terminus of the protein (Figure 4A). We also designed another construct in which we inserted a GSGS linker between the catalytic U-box domain of CHIP and BirA (CHIP-GSGS-BirA) (Figure 4A), under the assumption that providing more degrees of freedom for BirA may improve BirA-mediated site specific biotinylation of (-2)AP-Ub and thus leads to more efficient target protein biotinylation. CHIP-BirA or CHIP-GSGS-BirA were transiently expressed in HEK-293 cells for 24 h along with (-2)AP-Ub. As CHIP directs target proteins for proteasomal degradation, we treated cells with the proteasome inhibitor MG132 to preserve substrates for the duration of the experiment. Untreated cells served as control. Streptavidin immunoblotting of whole cell lysates showed that in the presence of MG132, expression of both CHIP-BirA and CHIP-GSGS-BirA resulted in a notable increase in biotinylation (Figure 4A). Interestingly, CHIP-GSGS-BirA showed more biotinylation compared to CHIP-BirA, suggesting that the linker improves the efficiency of substrate biotinylation, at least in the case of CHIP (Figure 4A). Since the structure of CHIP in complex with E2 is known(*61*), we designed a mutant of CHIP (I235D/F237D) that is deficient in binding to E2 as a negative control (Figure 4A). As expected, the biotinylation is notably lower in the condition where CHIP I235D/F237D transfected compared to CHIP WT.

**Figure 4:**
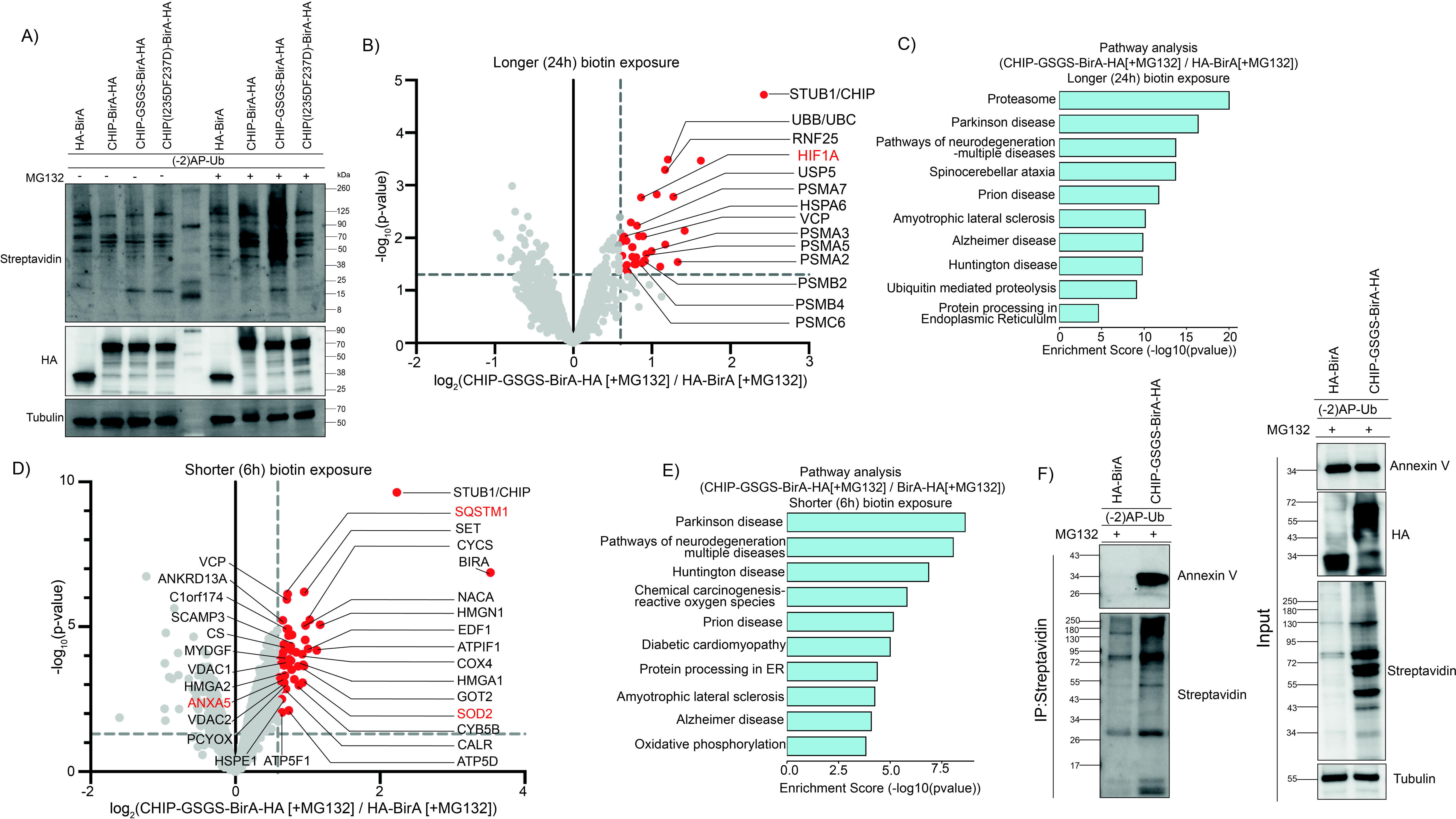
Employing Ub-POD to identify substrates of U-box ligase CHIP. A) HEK-293 cells transfected for 24 h with (-2)AP-Ub and indicated BirA constructs were treated with MG132 (10 µM) for 6 h. Cells were kept in biotin (100 µM) the whole time. Lysates were separated by SDS-PAGE and analyzed by immunoblotting. B) HEK-293 cells were transfected with (-2)AP-Ub and HA-BirA or CHIP-GSGS-BirA-HA as in A. Lysates were subjected to streptavidin IP and MS analysis. Volcano plot of proteins labeled by CHIP-GSGS-BirA-HA and HA-BirA in 24 h (n=3 biological replicates). Significantly altered proteins with p-value <0.05, log2FC >0.6 are colored as red circles and considered as potential CHIP substrates. Known substrate are highlighted in red. C) GO term analysis of significantly enriched candidate CHIP substrates. D) HEK-293 cells transfected for 24 h with (-2)AP-Ub and HA-BirA or CHIP-GSGS-BirA-HA were incubated with biotin (100 µM) and MG132 (10 µM) for 6 h. Lysates were subjected to streptavidin IP and MS analysis (n=3 biological replicates). Volcano plot of proteins labeled by CHIP-GSGS-BirA-HA and HA-BirA in 6 h (n=3 biological replicates). Significantly altered proteins are highlithed (p-value < 0.05, log2FC >0.6). Known CHIP substrates are marked in red. E) Bar graph representation of enriched GO terms for candidate CHIP substrates. F) HEK-293 cells transfected for 24 h with (-2)AP-Ub and HA-BirA or CHIP-GSGS-BirA-HA were treated with MG132 (10 µM) for 6 h. Cells were kept in biotin (100 µM) the whole time. Lysates were subjected to streptavidin IP followed by SDS-PAGE and immunoblotting. 5 % of lysates were used as input.

We chose CHIP-GSGS-BirA for the identification of potential substrates through proteomics. We performed the Ub-POD transfections as above and used expression of BirA alone as a control. Notably, cells in both conditions were treated with MG132. We did not induce proteotoxicity in this experiment, since we sought to identify substrates that get ubiquitinated by CHIP under basal conditions. Hypoxia inducible transcription factor, HIF1A, a well-known substrate of CHIP(*62*), was specifically identified in CHIP-GSGS-BirA expressing cells (Figure 4B, supplementary table 6). In addition, CHIP is reported to tag proteasomal subunits with polyubiquitin chains and target them to aggresomes(*63*). In agreement with this, several proteasomal subunit proteins were also identified as hits in the MS analysis (Figure 4B, supplementary table 6). Several neurodegenerative diseases, e.g. Parkinsons disease, Spinocerebellar ataxia, Alzheimer’s disease, Huntingtons disease (p<0.0001) were found enriched in GO term analysis in agreement with the known physiological role of CHIP (Figure 4C).

Since CHIP is involved in several different cellular processes apart from protein quality control(*64*), we wondered whether we could identify additional or a different set of substrates of CHIP by reducing the exposure of biotin in CHIP-GSGS-BirA expressing cells. Therefore, instead of exposing the cells to biotin immediately after transfection for 24 h, we added biotin to cells 24 h after transfection for 6 h along with MG132 (Figure 4D). Interestingly, two other known substrates of CHIP, SQSTM1 (also known as p62)(*65*) and SOD2(*66*) were identified (Figure 4D, Supplementary table 7) with this shorter biotin exposure. Notably, changes in the substrate identification profile depending on the duration of biotin pulse could be due to the different half-lives of various substrates of CHIP. GO annotation analysis of hits identified biotin treatment regimes, however, highlighted similar processes relevant to known functions of CHIP, such as pathways of neurodegeneration (p≤0.0001), and protein processing in the endoplasmic reticulum (p≤0.01) (Figure 4C and 4E).

Since we have used MG132 in both CHIP-GSGS-BirA as well as BirA expression conditions so far, the identified substrates may not necessarily be the ones that are strictly regulated through the proteasome (Figure 4B, 4D). Therefore. we next performed a comparison of Ub-POD CHIP in the presence or absence of MG132 (Supplementary figure 6 A-C, Supplementary table 8) to identify CHIP substrates targeted for proteasomal degradation. GO annotation analysis showed ubiquitin mediated proteolysis as the most significantly enriched pathway (p<0.0001) in presence of MG132 (Supplementary figure 6A-C). Previously known substrates such as SQSTM1(*65*), NPM1(*67*), PSMD4(*68*) and HDAC6(*69*) were identified in this Ub-POD analysis (Supplementary table 8). Applying Ub-POD to CHIP demonstrated that a flexible linker between BirA and the candidate Ub ligase can improve the identification of substrates and that biotin exposure times and stimuli may need to be optimized for a given Ub ligase to identify different sets of substrates.

Comparison of the putative CHIP substrates identified by Ub-POD to the ones previously identified through an orthogonal ubiquitin transfer (OUT) screen(*15*) revealed that among the 311 potential CHIP substrates identified by Ub-POD, 30 proteins overlapped with the 226 putative CHIP substrates found in the OUT screen (Supplementary Figure 6D; Supplementary table 9). Of these 30 overlapping proteins, only SQSTM1 and CTNNB1 were already known substrates of CHIP. However, two other well-known substrates of CHIP, SOD2 and HIF1, which were identified by Ub-POD, were not found in the OUT screen for CHIP.

Moreover, Annexin 5 (ANXA5), a member of the annexin superfamily of proteins, was identified by Ub-POD as well as by OUT screen (Figure 4D). Annexins are a family of cytosolic proteins that translocate to the plasma membrane in a calcium-dependent manner and thus regulates actin cytoskeleton dynamics(*70*). Intriguingly, a previous report found that ANXA5 is upregulated in CHIP KO cortical neurons compared to CHIP expressing neurons(*71*). Based on this, we hypothesized that ANXA5 is a possible substrate of CHIP ligase. In order to test this, we overexpressed either BirA or CHIP-GSGS-BirA in HEK-293 cells along with (-2)AP-Ub. Biotinylated proteins were enriched using streptavidin IP and probed by anti-ANXA5 antibody (Figure 4F). We observed specific enrichment of ANXA5 in condition where CHIP was expressed but not in control conditions indicating that ANXA5 is indeed a substrate of CHIP.

### Comparison of BioID and Ub-POD for E3 ligase substrate identification

Proximity labeling approaches such as BioID, TurboID and AirID have been used before to identify transient protein-protein interactions(*72–74*). These methods depend on a mutated form of BirA (BirA* which as R118G mutation) or other engineered biotin ligases fused to the protein of interest (POI) to release highly reactive AMP-Biotin into the surrounding solvent and label the “neighboring” macromolecules with biotin. Unlike BirA*, which is used in BioID, wildtype BirA employed in Ub-POD does not ligate biotin non-specifically to all the spatially proximal proteins but only attaches biotin to the (-2)AP-Ub that is in close proximity to E3. Nevertheless, we compared the suitability of BioID and Ub-POD for the identification of E3 ligase substrates.

Thereto, RAD18 was tagged at its N terminus with BirA*(*73*, *75*). BirA* alone or BirA* tagged RAD18 were expressed in HEK-293 cells. UV exposure and biotin treatment in the BioID experiment were followed exactly as for Ub-POD. Proteomics analysis showed that BioID-RAD18 identified numerous possible interacting and neighboring proteins of RAD18 (Figure 5A, Supplementary table 10). Functional annotation analysis of BioID-RAD18 hits revealed significant enrichment of GO categories related to spliceosome, basal transcription factors and homologous recombination (p<0.0001) (Figure 5B). Compared to 13 hits identified in RAD18 Ub-POD, BioID-RAD18 identified 320 hits (Figure 5C) providing a complex snapshot of RAD18’s many interacting and neighboring proteins. Surprisingly, we did not found any overlap between the hits identified by BioID and Ub-POD. Moreover the *bona fide* substrate of RAD18, PCNA was not identified in our BioID analysis (Figure 5A-C). To further explore the differences in BioID and Ub-POD, we also performed BioID analysis for CHIP. CHIP was tagged with BirA* at its C terminal end (CHIP-BirA*). MG132 and biotin treatment were kept the same for BioID-CHIP as for CHIP Ub-POD (Figure 4B). GO pathway analysis of the hits identified by CHIP-BirA* compared to BirA* unveiled pathways related to various cellular roles of CHIP with significant p values (p<0.0001) (Figure 5D-E). Among the identified CHIP-BirA* hits were heat shock proteins, proteasomal subunit proteins and a number of known CHIP substrates (SOD2, SOD1, CDK4, ANXA5) (Figure 5D, Supplementary table 11). Comparison of all 1036 hits identified in BioID-CHIP (Figure 5D) with the 43 hits identified by CHIP Ub-POD (Figure 4B, 4D) showed 17 common hits between these two datasets (Figure 5F, Supplementary table 12). These results indicated that, though both BioID and Ub-POD are dependent on proximity-based biotinylation of target proteins, Ub-POD identifies far fewer number of hits compared to BioID likely because it is specific to the Ub pathway proteins and substrates.

**Figure 5:**
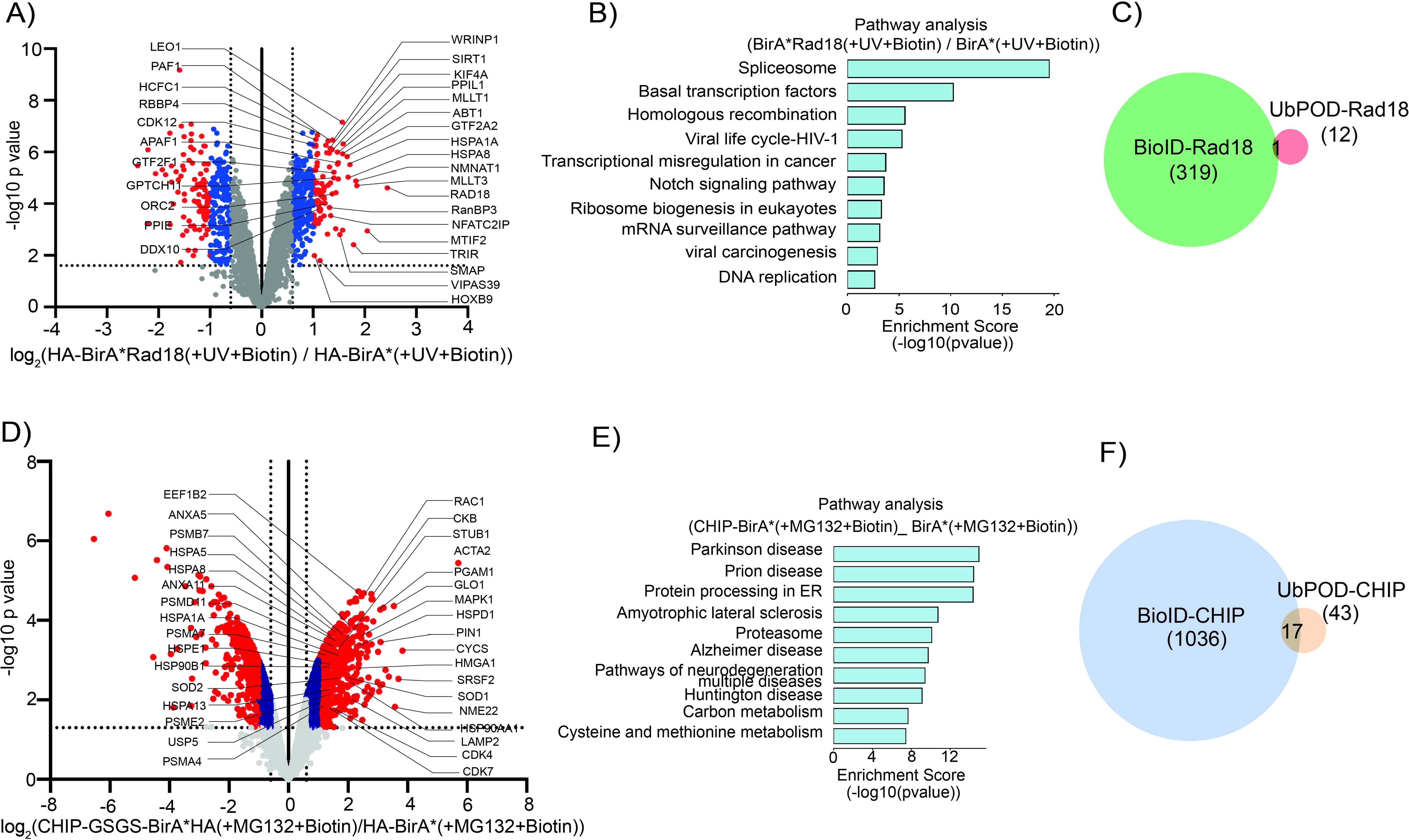
BioID of RAD18 and CHIP. A) BioID analysis of Rad18. HEK-293 cells transfected for 24 h with BirA* or BirA*-Rad18 were exposed to UV for 6 h. Cells were kept in 100 µM biotin the whole time. Lysates were subjected to streptavidin IP followed by MS. Volcano plot of proteins labeled by BirA*-Rad18 and BirA*. Significantly altered proteins are marked in blue (p-value ≤0.05, 0.6≤ log2FC <1) and red (p-value ≤0.05, 1< log2FC). B) Bar graph depicting significantly enriched GO terms of hits from A. C) Overlap between hits identified in BioID-Rad18 and RAD18 Ub-POD. D) BioID analysis of CHIP. HEK-293 cells transfected for 24 h with BirA* or CHIP-GSGS-BirA* were incubated with MG132 (10uM) for 6 hours. Cells were kept in 100 µM biotin the whole time. Streptavidin IPs were performed with lysates followed by MS analysis. Volcano plot of proteins labeled by CHIP-GSGS-BirA* and BirA*. Significant hits are labeled in blue (p-value ≤0.05, 0.6≤ log2FC <1) and red (p-value ≤0.05, 1< log2FC). E) Bar graph representation of significantly enriched GO terms with hits identified in D. F) Overlap between hits identified in BioID-CHIP and CHIP Ub-POD.

## Discussion

Here, we developed a new approach to effectively label the substrates of a given ubiquitin ligase with biotin directly in cells and identify them using quantitative MS. It is worth mentioning that our approach has important differences from the traditional proximity labeling approaches such as BioID and TurboID, which have been successfully employed in a number of studies to identify interacting partners of a POI(*72*, *73*). The BioID method involves the expression of the POI tagged with a mutant of BirA, called BirA* or BioID, which bears the mutation R118G in the active site of the enzyme. BirA* releases highly reactive biotin-AMP conjugate into solvent which can label any protein within the spherical radius of ∼10nm, thus providing a clear picture of the interactors and the environment of the POI. However, BioID does not differentiate between the direct interactor and any other protein within the vicinity of the POI. In the context of E3 ligases, BioID would also not be able to differentiate between the substrate of the ligase and an interactor of the ligase. These predictions are reflected in our data when we performed BioID of Ub ligases RAD18, CHIP and compared the identified hits with those of Ub-POD (Figure 5). We introduced two essential tweaks into the principle of BioID to make it selective for ubiquitin (Ub). Firstly, we fused the BirA WT, which can only efficiently biotinylate the lysine present in the acceptor peptide (AP) or Avi-tag, to the E3 ligase of interest. Secondly, we attached a variant of AP-tag ((-2)AP) to the N-terminus of Ub to be selectively biotinylated by BirA WT in a proximity-dependent manner in the course of substrate ubiquitination (Figure 1A-B). We hypothesized that Ub-POD is also sensitive to the location of the catalytic domain in the E3 ligase, since according to the catalytic intermediate complex structures(*19*, *23*, *24*, *76*) (Supplementary Figure 1A, 1B), fusion of BirA to the opposite end of the protein relative to the catalytic domain likely places it too far from the (-2)AP-Ub∼E2. For efficient biotinylation, we fused BirA to the N-terminus of RAD18, which possesses a RING domain at the N-terminus, and to the C-terminus of CHIP, which contains U-box domain at the C-terminus. When we fused the tag at the non-catalytic ends of RAD18 and CHIP, we still observed higher biotinylation of cellular proteins compared to the background caused by BirA alone (Supplementary Figure 7A-B). However, we have not performed a comparison of substrates identified with different tag placements, an analysis which is required to test if the BirA tag placement plays a significant role in the Ub-POD of E3 ligases. We predict that is likely to be case-specific. Intriguingly, we also observed that the linker between the BirA and the E3 ligase also plays an important role in determining the efficiency of the Ub-POD. By increasing the linker length in the case of CHIP, we were able to increase the biotinylation (Figure 4). Furthermore, we discovered that both the timing of biotin addition to the cells relative to the expression of the BirA-tagged ligase and the duration of biotin exposure can also be important parameters that can be varied, depending on the E3 ligase of interest (as is the case of CHIP, Figure 4), to allow the detection of different sets of substrates for a given E3 ligase.

The primary condition for the success of Ub-POD for a given ubiquitin ligase is that the ligase-mediated substrate ubiquitination should occur and be preserved until isolation in the cell line of choice and under the conditions employed. Use of appropriate triggers, such as UV for RAD18, should be considered for activating ligases. For preserving ubiquitination and to prevent Ub recycling, the use of MG132 and/or PR619 or other similar agents should be considered. For substrate identification through proteomics, we primarily expressed BirA alone without the candidate ligase fusion as a control. The successful use of BirA as a control for multiple ligases here also implies that it could serve as an efficient control, in principle, for other RING or U-Box E3 ligases. Where known, point mutants of ligases lacking the catalytic activity can also be used as controls. Our data indicates that for ligases known to homodimerize, activity-deficient mutants might not act as efficient controls in the Ub-POD MS experiments (Supplementary Figure 2).

Several available methods can also be employed to identify the substrates of ubiquitin ligases(*6*, *7*). Although the gold standard methods, such as diGly and GPS profiling (specific for degradative ubiquitination), are very effective in the identification of ligase substrates, they are resource intensive and require acquired expertise to employ(*8–10*). Simpler methods similar to Ub-POD that trap the substrate of ubiquitin ligases also exist, such as UBAIT or TULIP, which depend on the fusion of Ub to the ligase to covalently trap the ligase-substrate complex(*11*, *13*). These methods depend on the ability of the ligase to use the Ub-ligase fusion, which is quite bulky compared to Ub alone, in the Ub-cascade reaction. Any laboratory with access to basic tissue culture and a mass spectrometry facility would be able to employ Ub-POD. Moreover, the (-2)AP-tag on Ub is very small compared to the Ub-ligase fusion employed in UBAIT and related approaches. Ub-POD also enables the isolation of potential substrates in harsh denaturing conditions which extracts proteins residing in typically insoluble cellular fractions and also limits the number of false positive hits. Finally, Ub-POD provides a unique opportunity to visualize the cellular localization of ubiquitination by a ligase of interest, offering a peek into the cellular function of the ligase.

In the current study, we demonstrated the use of Ub-POD on two classes of E3 ligases, RING and U-box. RING-type E3 ligases comprise the majority of ubiquitin ligases in humans and the principles of Ub-POD can be applied for substrate identification of all of ∼600 RING E3 ligases.

Although not part of this study, we predict that Ub-POD can be employed for HECT or RBR E3 ligases, which also operate on principles of complexing with Ub before transfer of Ub to the substrate. Ubiquitin-like molecules (UBLs) such as SUMO, NEDD8, FAT10, ISG15 and UFM1 are also conjugated to proteins in a similar fashion to Ub, using a near-identical cascade of enzymatic reactions(*77*). By tagging the respective ligases with BirA and UBL with (-2)AP-tag, Ub-POD can in principle be adapted to other UBLs as well. In light of the demonstrated applications and possible other applications discussed here, Ub-POD likely will play an important role in future ubiquitin research.

---

**Supplementary Figure 1:**
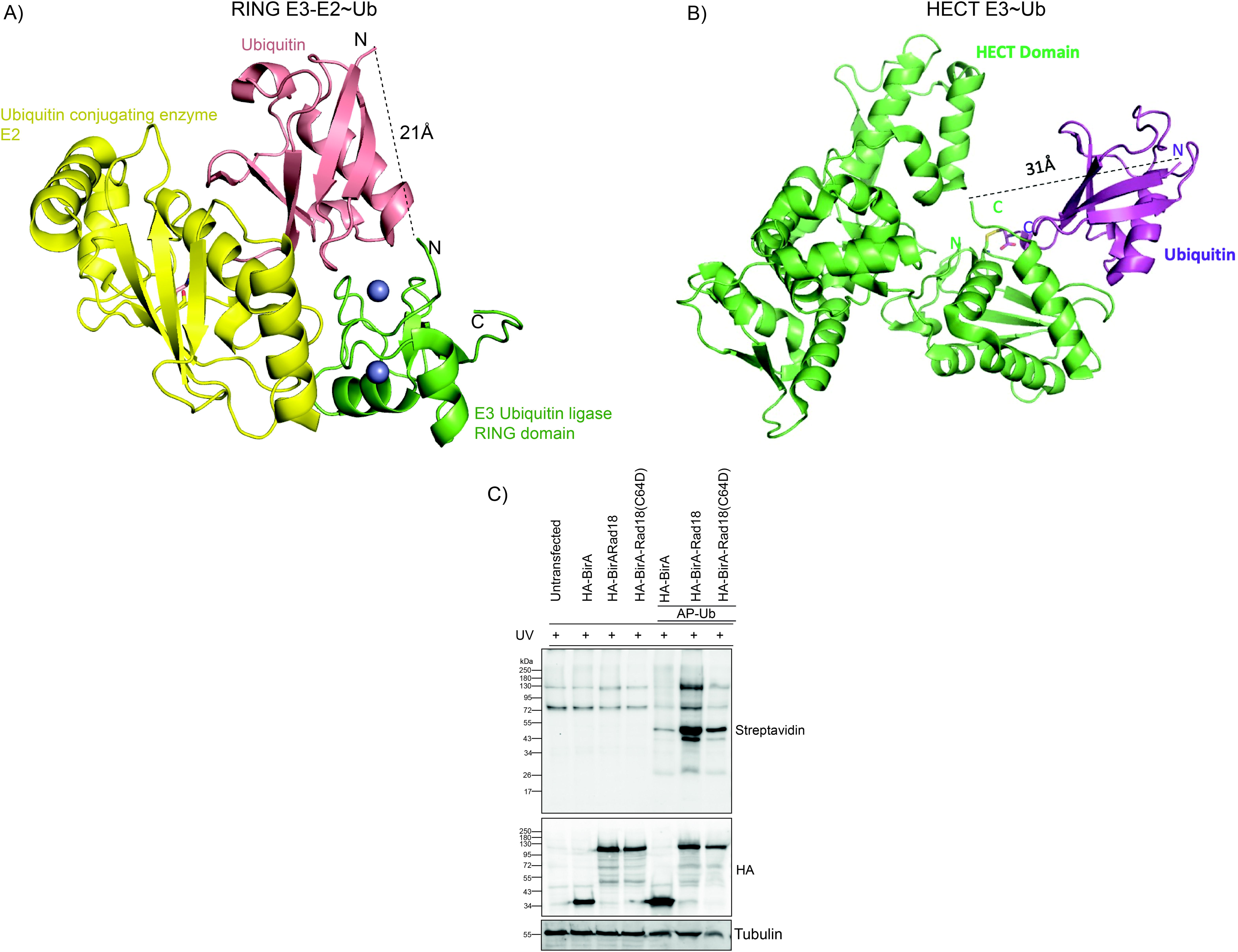
A) Structural model of RING E3 ligase (green) in complex with ubiquitin conjugating enzyme E2 (yellow) and ubiquitin (red). The distance between the N-terminus of Ub and the catalytic RING domain is shown. B) Structural model of HECT-E3-ligase (green) in complex with Ub (purple) showing the distance between the catalytic HECT domain and the N-terminus of Ub. C) HEK-293 cells transfected for 24 h with the indicated constructs were exposed to UV for 6 h. Cells were kept in 100 µM biotin the whole time. Lysates were subjected to SDS-PAGE and streptavidin immunoblotting.

**Supplementary Figure 2:**
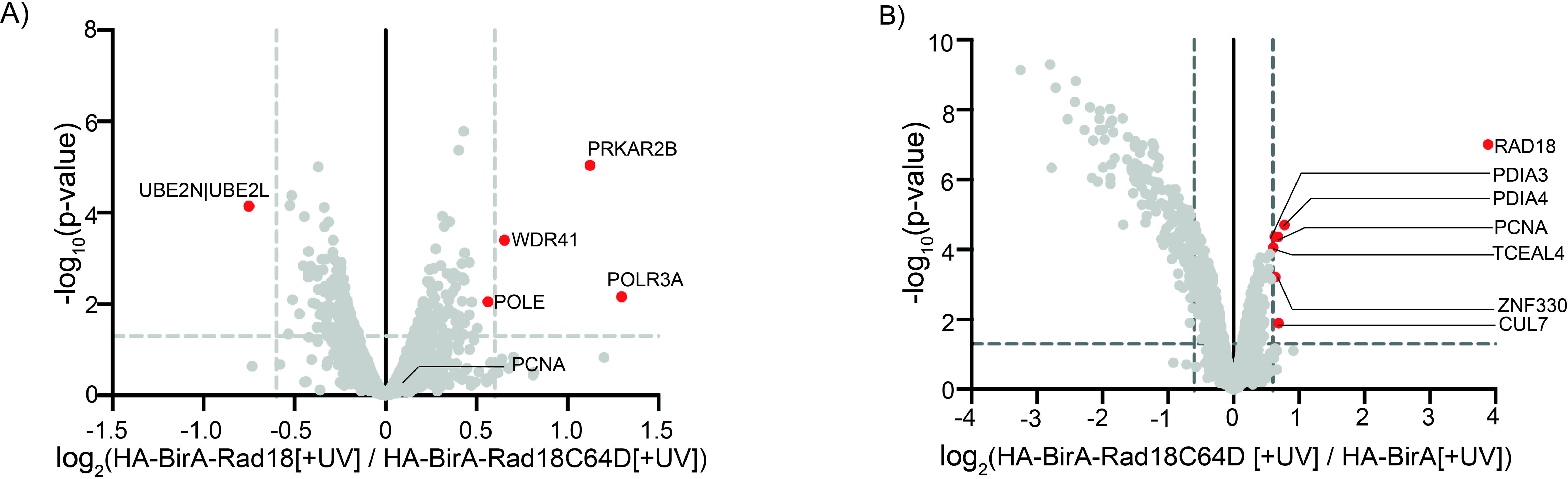
A) Volcano plot depicting differentially enriched biotinylated proteins from streptavidin IP samples of HA-BirA-Rad18 WT and HA-BirA-Rad18 C64D expressing HEK-293 cells treated with biotin and UV. Significantly altered proteins are labeled in red (p-value ≤0.05, log2FC ≥0.6). B) Volcano plot of proteins labled HA-BirA-Rad18 C64D and HA-BirA in cells treated with biotin and UV. Significantly altered proteins (p-value ≤0.05, log2FC ≥0.6) are highlighted in red.

**Supplementary figure 3:**
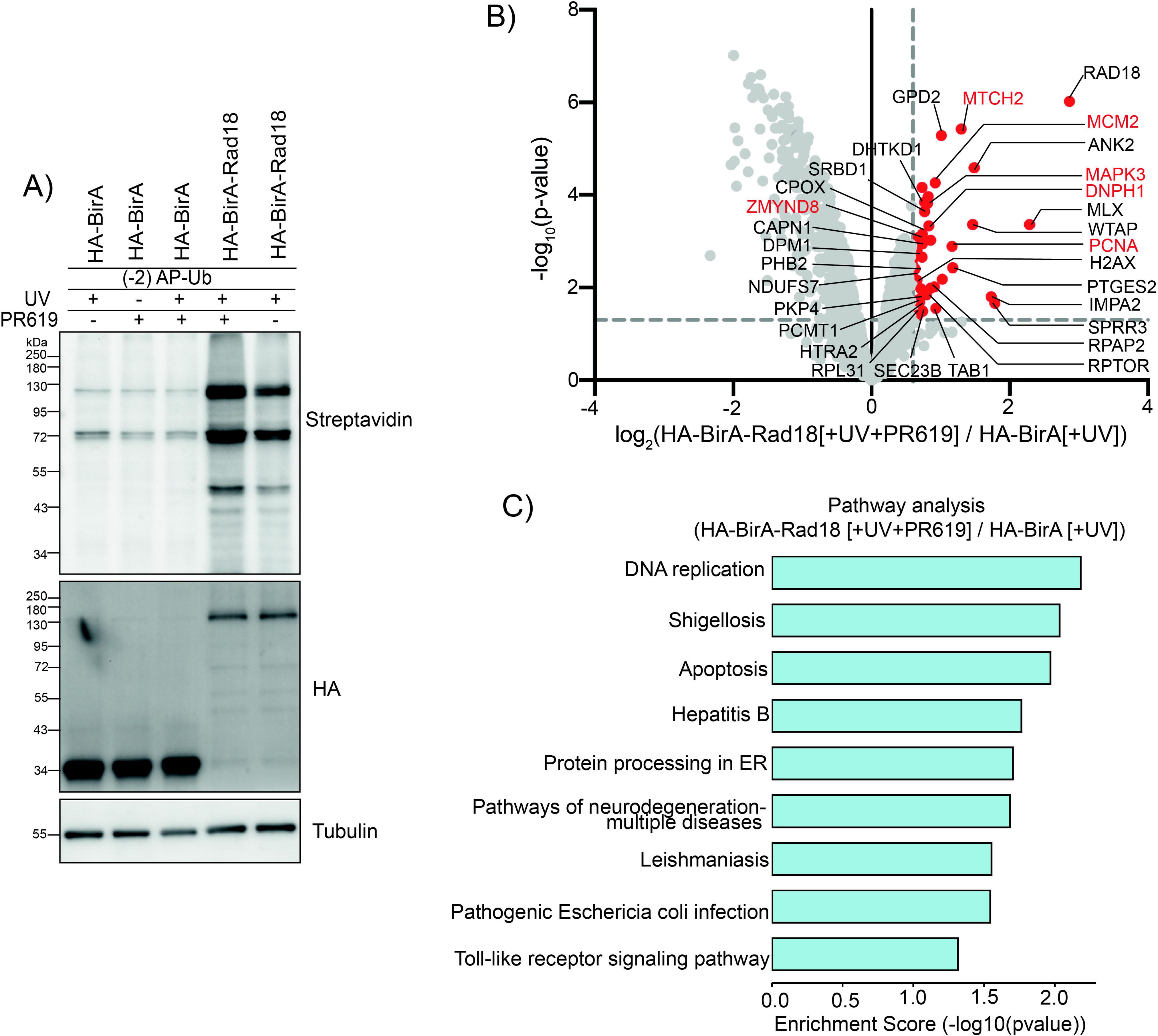
A) HEK-293 cells transfected for 24 h with (-2)AP-Ub and HA-BirA or HA-BirA-Rad18 followed by treatment with UV and PR619 (10uM) as indicated. Cells were kept in biotin (100 µM) the whole time. Lysates were subjected to SDS-PAGE and streptavidin immunoblotting. B) MS analysis of streptavidin IP samples (n=3 biological replicates) of HA-BirA and HA-BirA-Rad18 transfected HEK-293 cells treated as described in A. Volcano plot of proteins labeled by HA-BirA and HA-BirA-Rad18. Significantly altered proteins (p-value <0.05, log2FC >0.6) are labelled in red. C) GO term enrichment analysis of identified hits are shown in bar graph.

**Supplementary Figure 4:**
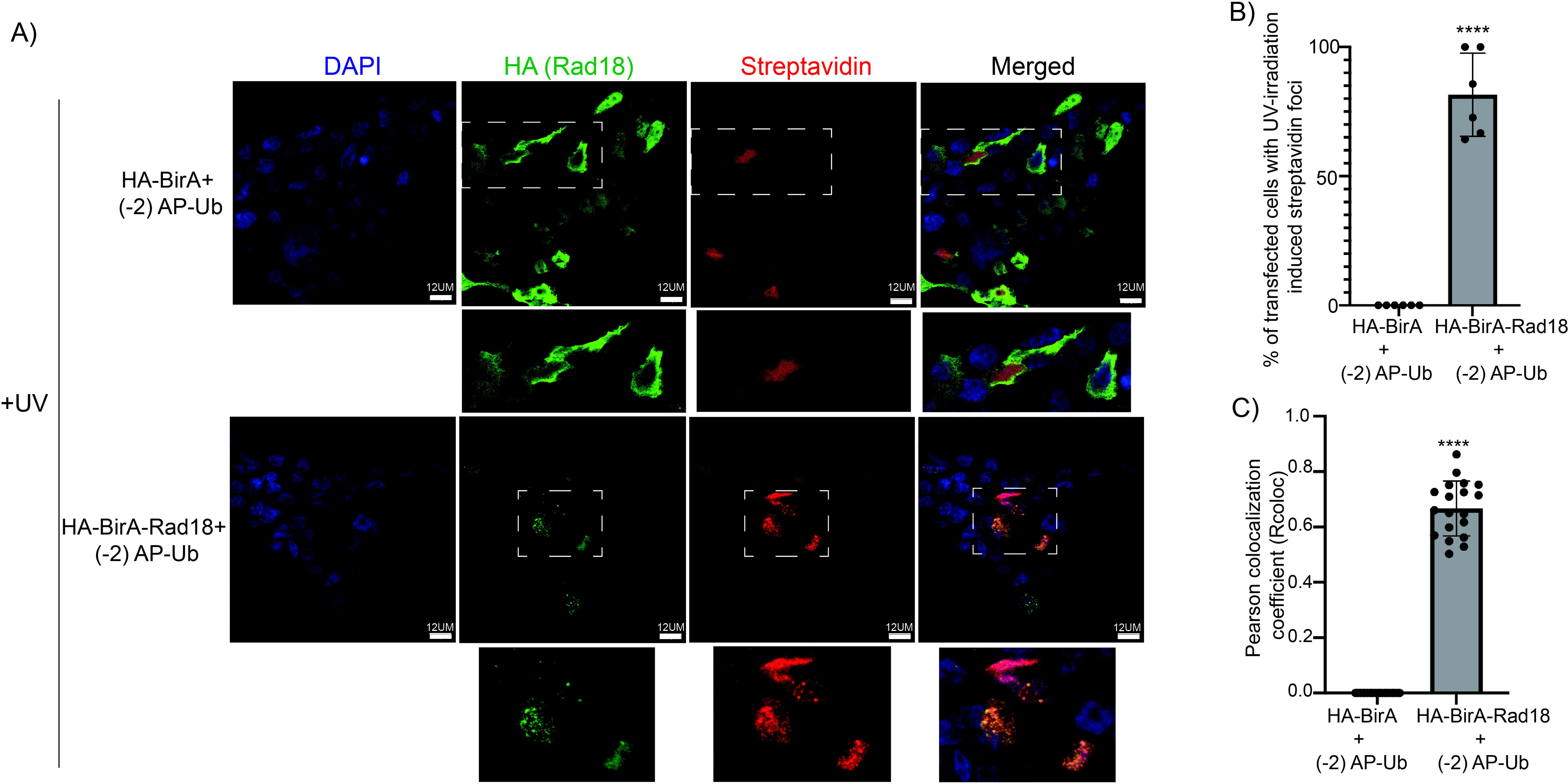
A) HEK-293 cells were transfected and treated as in B followed by fixation and immunostaining with anti-HA (green) and anti-streptavidin (red) antibodies. DAPI was used to stain the nucleus (blue). Scale bar 12µm. B) Quantification of experiment shown in Figure E. Percentage of transfected cells with UV irradiation-induced foci were counted for HA-BirA and HA-BirA-RAD18 (****p<0.0001; n= 6 different fields). C) Pearson correlation coefficient was measured to depict colocalization between red and green channels for HA-BirA and HA-BirA-RAD18 and represented as a bar graph (****p<0.0001; n=20 cells analysed for each condition from different fields).

**Supplementary Figure 5:**
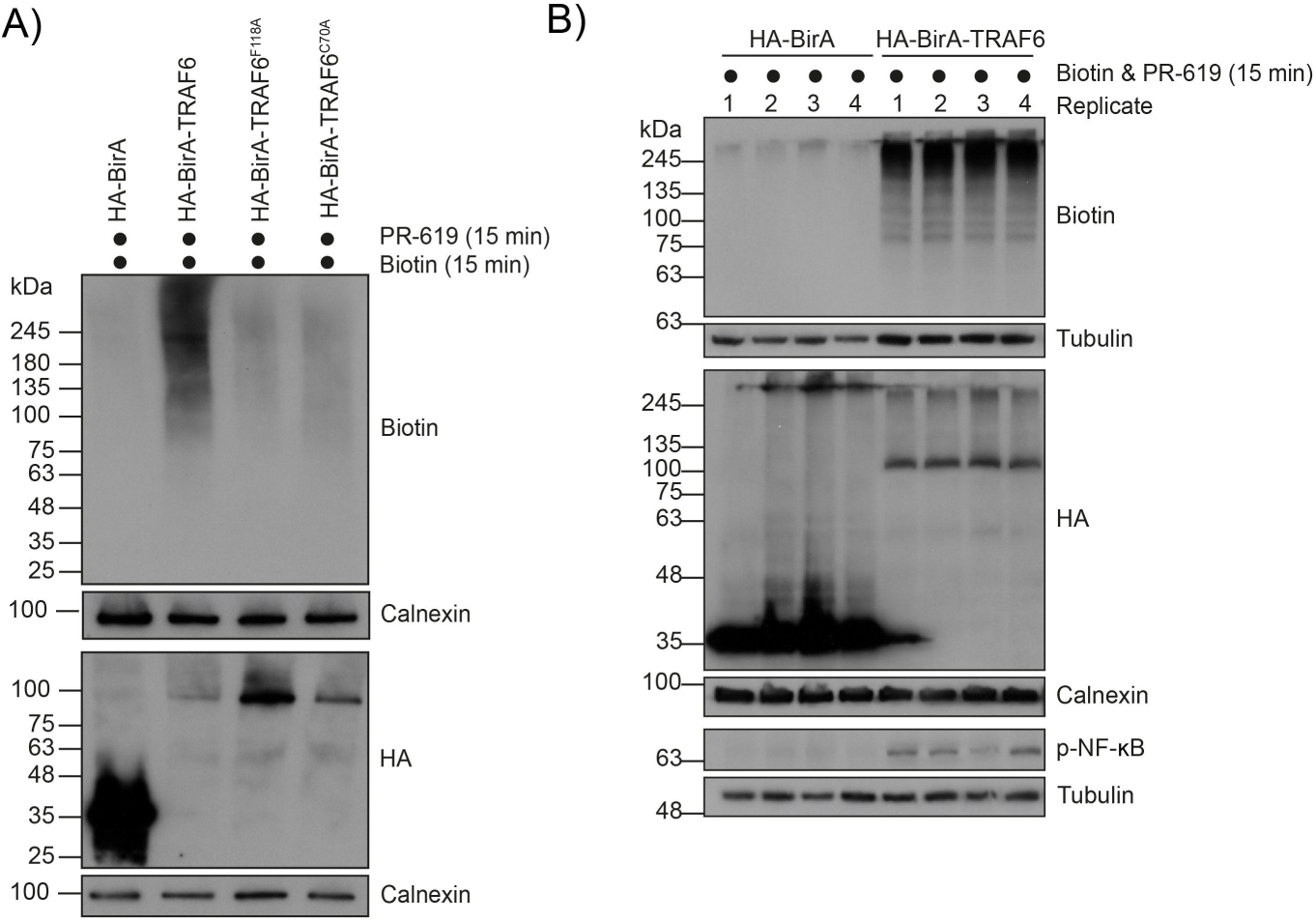
A) Biotinylation specificity control for TRAF6. After 24 h transfection, cells transfected with indicated constructs were treated with 100 µM biotin and 10 µM PR-619 for 15 min. Lysates were separated by SDS-PAGE and analyzed by immunoblotting. B) Aliquots of MS sample from Figure 3C (4 biological replicates) were analyzed by SDS-PAGE and immunoblotting.

**Supplementary Figure 6.**
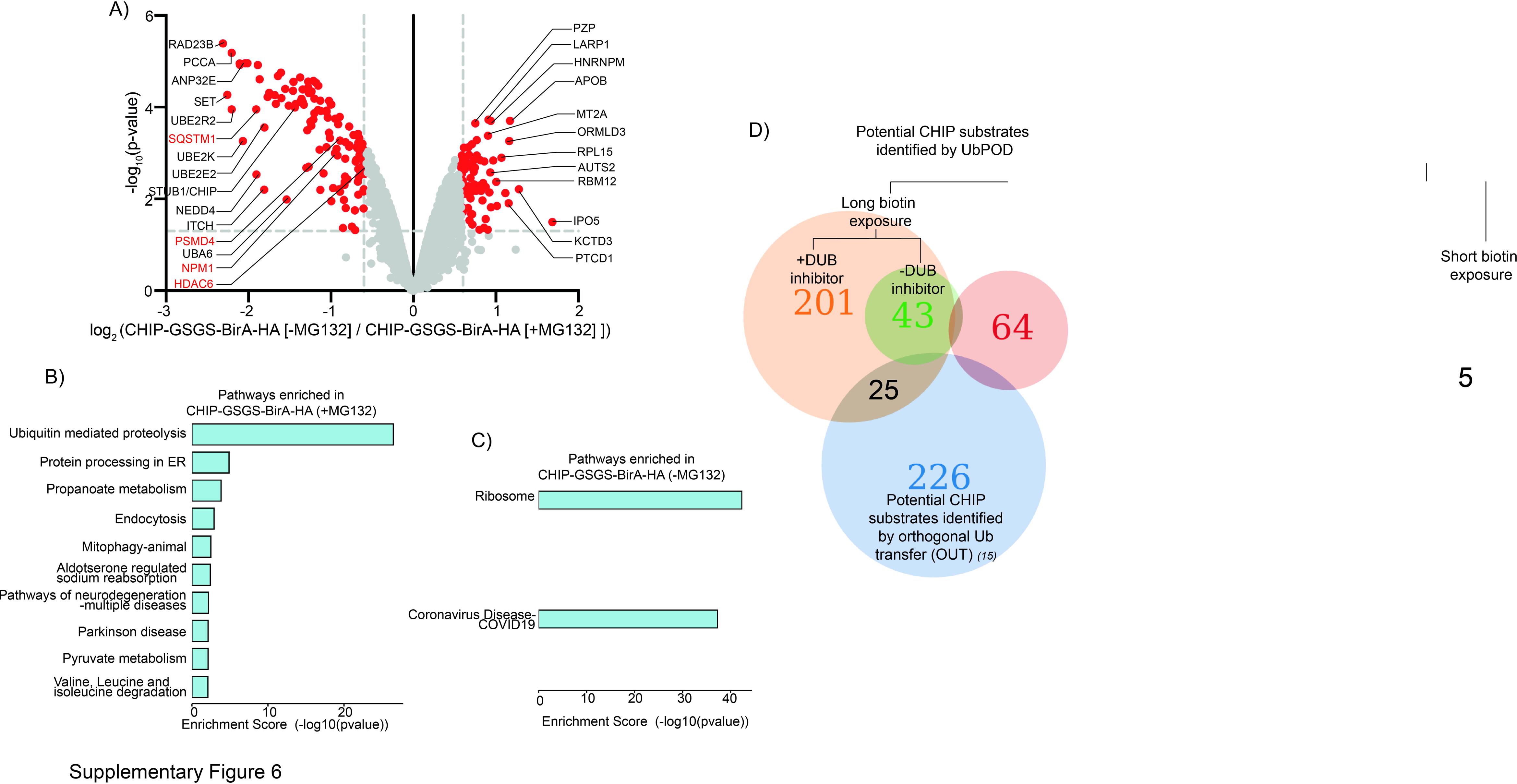
A) MS analysis of streptavidin IPed samples from CHIP-GSGS-BirA-HA expressing HEK-293 cells grown in the presence or absence of MG132 (n=3 biological replicates). Volcano plot of proteins labeled by the CHIP in MG132 treated and untreated cells. Significantly altered proteins are coloured in red (p-value <0.05, -0.6≥ log2FC ≤0.6). Known substrates are highlighted in red. B and C) GO term enrichment analysis of hits identified by CHIP-GSGS-BirA-HA in the presence MG132 (B) and absence of MG132 (C). Overerepresented pathways are shown in bar graph. D) Venn diagram showing overlap of identified proteins in proteomics analysis of different CHIP Ub-POD experiments and CHIP-OUT screen (*15*).

**Supplementary Figure 7:**
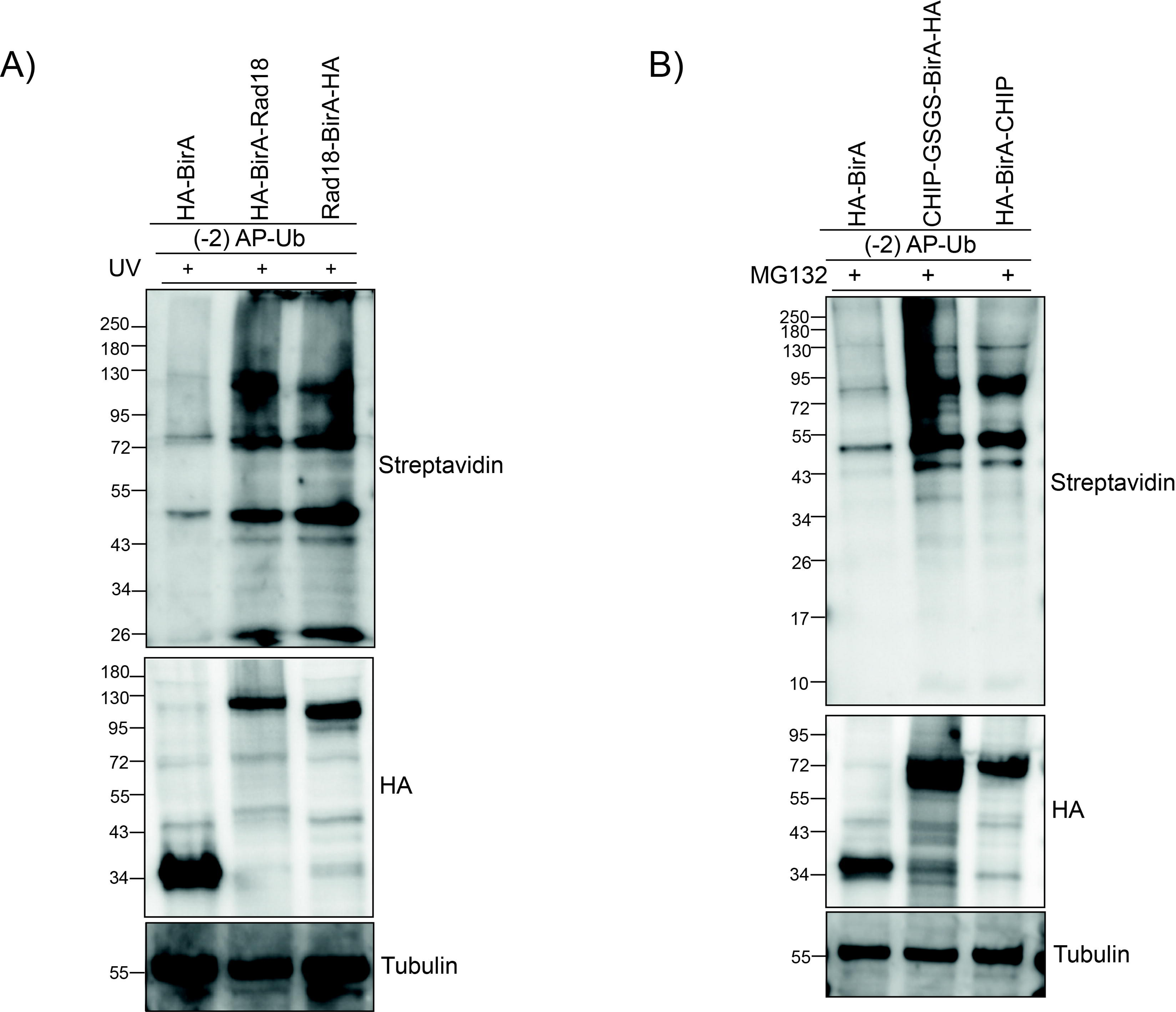
A) HEK-293 cells transfected with (-2)AP-Ub and HA-BirA*, HA-BirA*-RAD18. or RAD18-BirA*-HA were incubated with biotin and exposed to UV for 6 h. Lysates were subjected to SDS-PAGE and streptavidin immunoblotting. B) HEK-293 cells transfected with (-2)AP-Ub and CHIP-GSGS-BirA-HA or HA-BirA-CHIP were incubated with biotin and MG132. Lysates were subjected to SDS-PAGE and streptavidin immunoblotting.

## Supplementary tables

**Table S1:** Proteomics of HA-BirA-RAD18 [+UV] versus HA-BirA-RAD18 [-UV]. Related to Figure 1D.

**Table S2:** Proteomics of HA-BirA-RAD18 [+UV] versus HA-BirA [+UV], HA-BirA-RAD18 [+UV] versus HA-BirA RAD18 C64D [+UV] and HA-BirA-RAD18 C64D [+UV] / HA-BirA [+UV]. Related to Figure S2C, 2C and S2B.

**Table S3:** Proteomics of HA-BirA-RAD18 [+UV+PR619] versus HA-BirA [+UV]. Related to Figure S3B.

**Table S4:** Proteomics of BirA-TRAF6 versus BirA. Related to Figure 3C.

**Table S5:** Hits of BirA-TRAF6 versus BirA. Related to Figure 3C.

**Table S6:** Proteomics of CHIP-GSGS-BirA-HA [+MG132] versus HA-BirA [+MG132] with 24 h biotin incubation. Related to Figure 4B.

**Table S7:** Proteomics of CHIP-GSGS-BirA-HA[+MG132] versus HA-BirA[+MG132] with 6 h biotin incubation. Related to Figure 4D.

**Table S8:** Proteomics of CHIP-GSGS-BirA-HA[+MG132] versus CHIP-GSGS-BirA-HA[-MG132]. Related to Figure S6D.

**Table S9:** Common CHIP substrates identified by CHIP Ub-POD and CHIP-OUT (15). Related to Figure S6G.

**Table S10:** Proteomics of HA-BirA*RAD18 (+UV) versus HA-BirA* (+UV). Related to Figure 5A.

**Table S11:** Proteomics of CHIP-GSGS-BirA*HA (+MG132) versus HA-BirA* (+MG132). Related to Figure 5D.

**Table S12:** Common CHIP substrates identified by BioID-CHIP and CHIP Ub-POD. Related to Figure 5.

## Acknowledgements

We thank the EMBL Mass spectrometry core facility, especially Mandy Rettel for the data acquisition and analysis, Martin Pelosse for training and maintenance of the EMBL eukaryotic expression facility, This work used the platforms of the Grenoble Instruct-ERIC center (ISBG; UAR 3518 CNRS-CEA-UGA-EMBL) within the Grenoble Partnership for Structural Biology (PSB), supported by FRISBI (ANR-10-INBS-0005-02) and GRAL, financed within the University Grenoble Alpes graduate school (Écoles Universitaires de Recherche) CBH-EUR-GS (ANR-17-EURE-0003). SB was supported by a grant from the French Agence Nationale de la Recherche (ANR-21-CE11-0013). UM was supported by EMBO Postdoctoral Fellowship (EMBO-ALTF-185-2021). CB was supported by the Deutsche Forschungsgemeinschaft (DFG, German Research Foundation) within the frameworks of the Munich Cluster for Systems Neurology (EXC 2145 SyNergy – ID 390857198 (C.B.) and the Collaborative Research Center (CRC) 1177 (ID 259130777).

## Author Contributions

UM performed Ub-POD experiments with RAD18 and CHIP. SL performed Ub-POD experiments with TRAF6. SG assisted in initial Ub-POD experiments. FS analyzed the mass spectrometry data. CB conceptualized Ub-POD on TRAF6 and supervised SL. SB conceptualized the study and supervised UM. SL, CB, UM, and SB wrote the manuscript.

## Materials and methods

### Cell culture

Human embryonic kidney epithelial cell line 293 [HEK-293] (ATCC CRL-1573) was cultured in Dulbecco’s Modified Eagle Medium (DMEM), supplemented with 10% (*v*/*v*) heat-inactivated Fetal Bovine Serum (FBS; Gibco™) within humidified 5% CO_2_ incubator at 37°C. The cell line used in this study was free from mycoplasma contamination based on PCR detection.

### Reagents and antibodies

All reagents used in this study are listed in Table 1 and reconstituted according to manufacturers’ instructions. Polyclonal and monoclonal antibodies used for this study are listed in Table 2 and were used according to the manufacturers’ recommended dilutions.

**Table 1:**
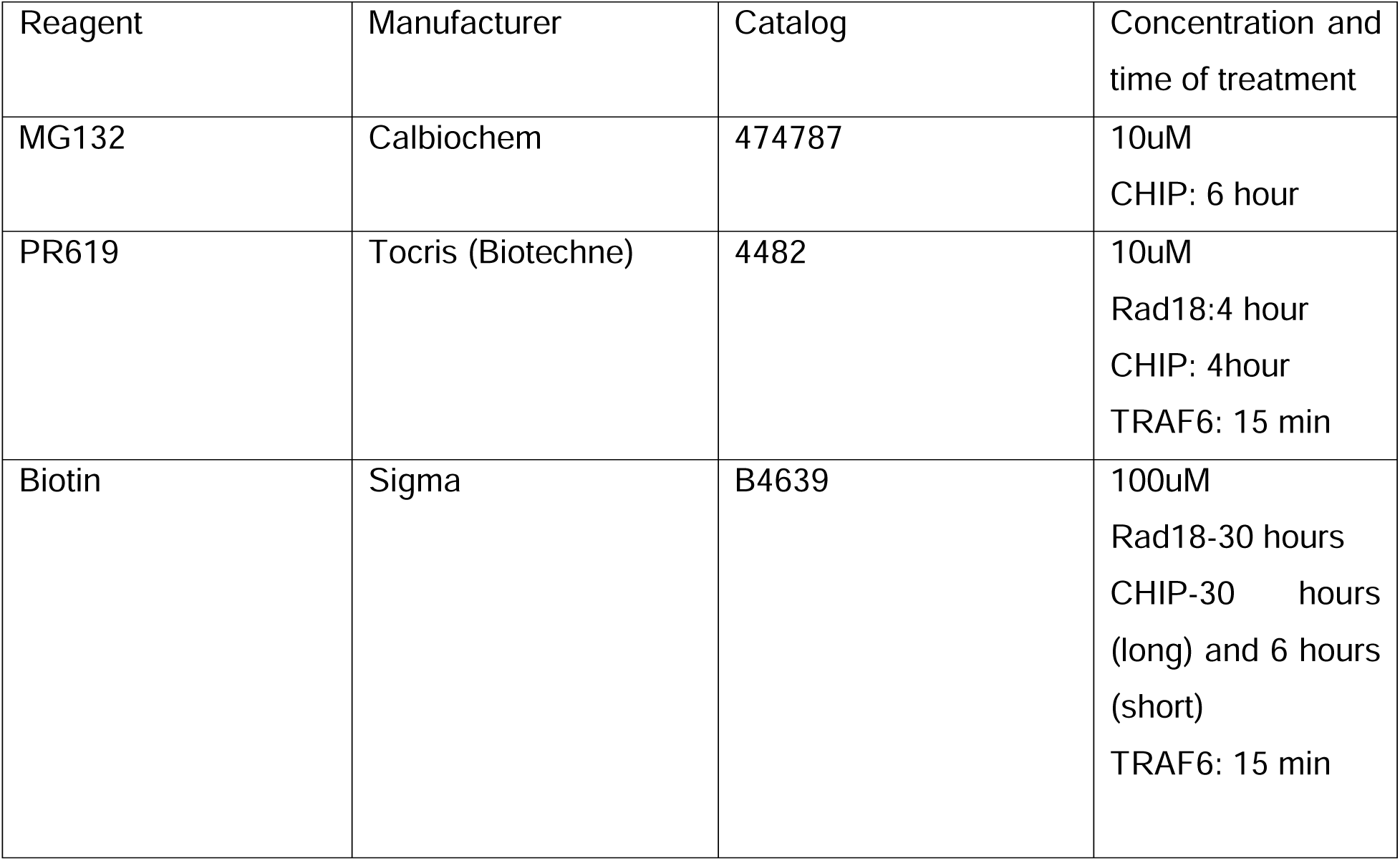
List of reagents.

**Table 2:**
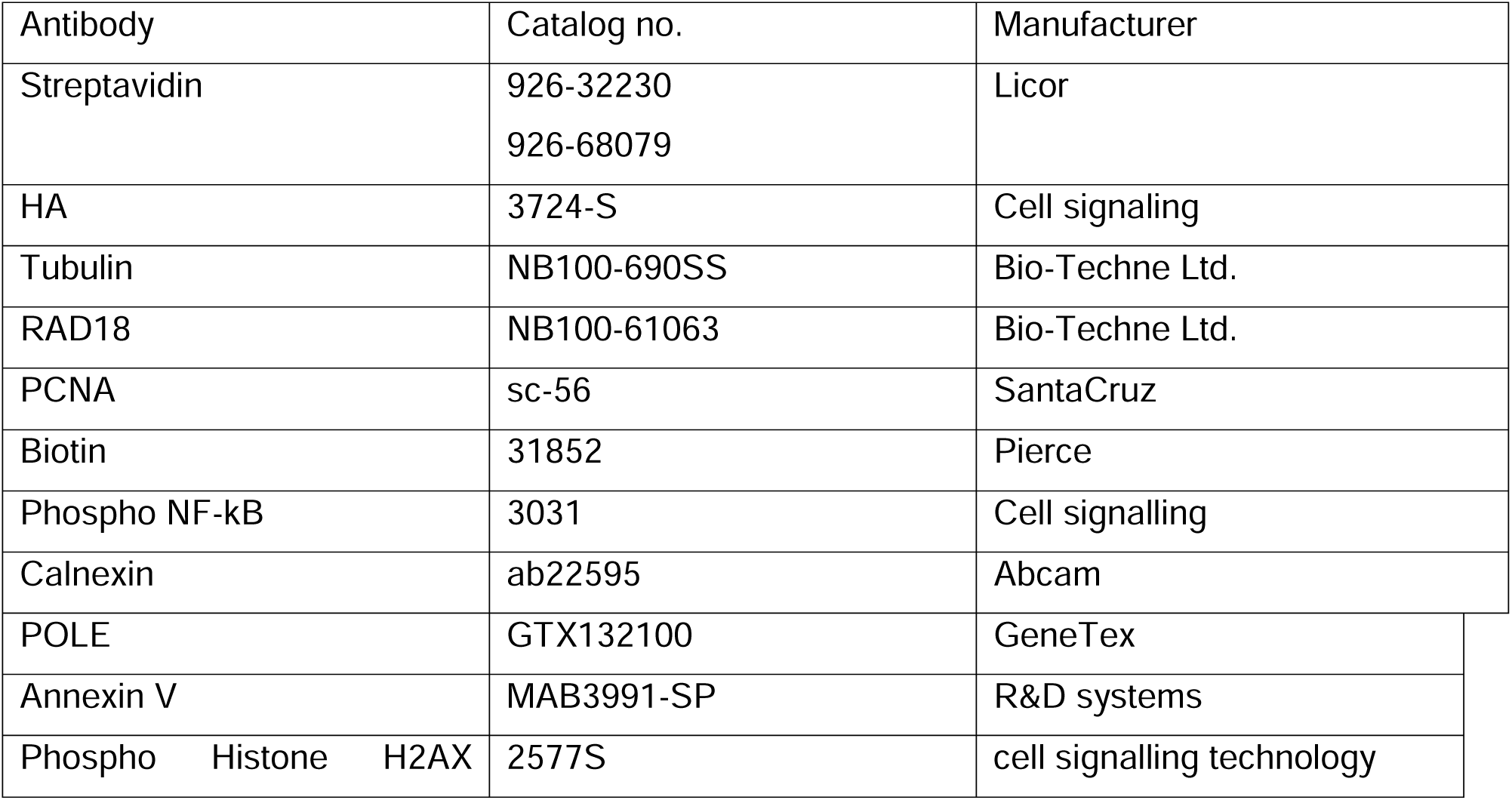
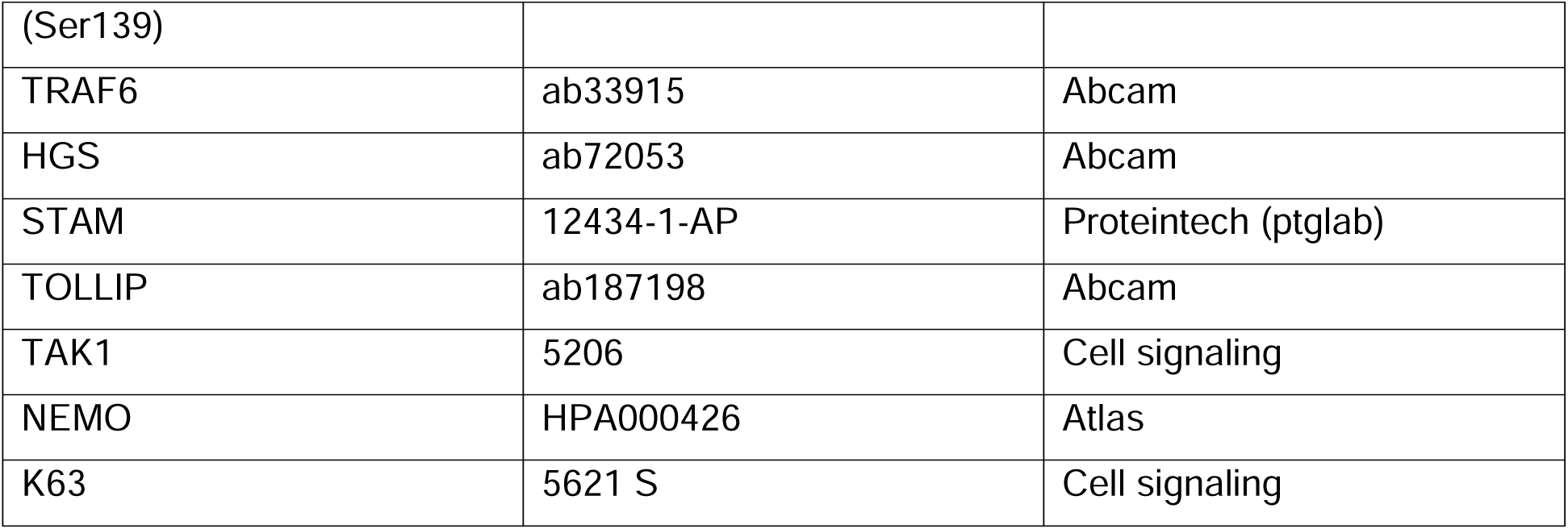
List of antibodies.

### Cloning

In order to tag the catalytic end of the candidate ligases, two vectors containing wild type E.coli biotin ligase enzyme BirA tagged with HA [HA-BirA for tagging candidate ligases with N terminal catalytic domain and BirA-HA for tagging candidate ligases with C terminal catalytic domain] were made, in which the candidate E3 ligases can be cloned based on the position of the catalytic domain. Full-length RAD18, CHIP and TRAF6 was incorporated into required BirA vector by sequence and ligation independent cloning. A GSGS linker was placed in between the catalytic end of candidate ligases and BirA by site directed mutagenesis (SDM). Mutant ligase constructs (either catalytic cysteine mutants or E2-E3 interaction interface mutants) were made also by SDM. Full length Ubiquitin was tagged at its N terminus with an avi tag (AP-Ub). To make truncated AP-Ub constructs (-2)AP-Ub and AP(-3)-Ub, 2 and 3 amino acids from N terminus and C terminus of the Avi tag were truncated, respectively. The list of constructs used in this study are presented in the table 3

**Table 3:**
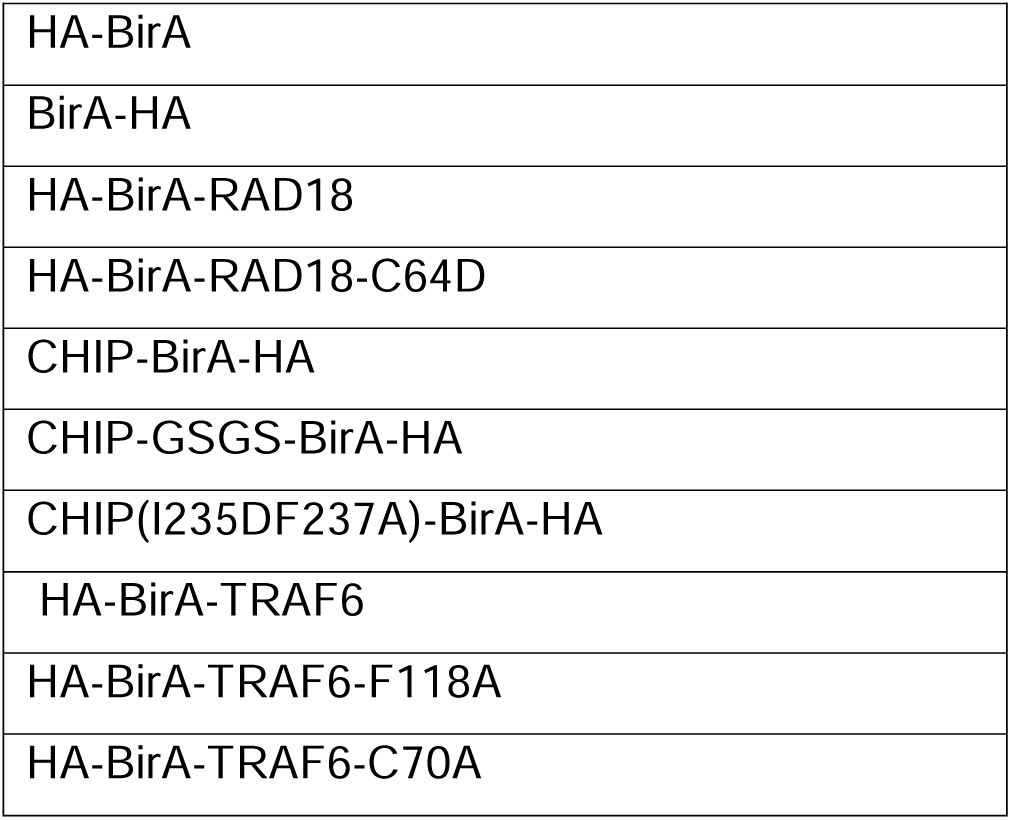
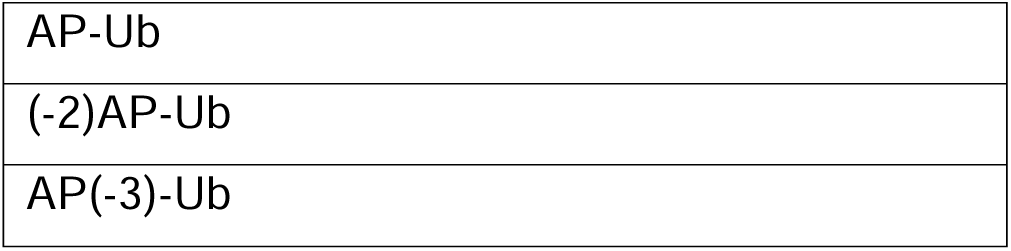
List of constructs.

### Transient transfection

All plasmids were transfected in HEK-293 cells with Polyethyleneimine (PEI) (Invitrogen) according to the manufacturer’s instructions. Briefly, the cells were seeded the day before transfection. Next day, equal amount of Avi-tagged Ub and BirA-tagged candidate ligase constructs were mixed with PEI in a reduced serum medium, the transfection mix was incubated at room temperature for at least 20 minutes before adding dropwise to the cells. The transfected cells were incubated overnight at 37°C, 5% CO_2_.

### Biotin labeling and induction of target protein ubiquitination

Different strategies were taken to induce target protein ubiquitination by the candidate ligases. For RAD18, UV exposure was used as an inducer of DNA damage. For CHIP, the proteasome inhibitor, MG132 was used as an inducer. For TRAF6 no inducer was necessary, since overexpression itself resulted in dimerization and thus activation of E3 ligase activity.

### Biotin labelling and UV exposure for RAD18

Cells were treated with biotin (100uM) once during transfection. After 24 hours of transfection, the cells were washed in 1X PBS and exposed to UV (10 J/m2) in a Stratalinker UV crosslinker oven (Stratagene).After UV exposure, PBS was replaced with biotin (100uM) supplemented complete media and the cells were further incubated for 6 hours of at 37°C, 5% CO_2_ incubator.

### Biotin labelling and MG132 treatment for CHIP

We took two different biotin labelling regimes for CHIP a) shorter biotin exposure: biotin treatment (figure 4D, 4E) was done only during the inducer (MG132) treatment, 24 hours after transfection. Briefly, the transfection media was removed and replaced with medium supplemented with 100 uM biotin (dissolved in DMEM) and 10uM MG132 and incubated at 37°C, 5% CO_2_ incubator for 6 hours. b) longer biotin exposure: biotin treatment was done once during transfection, as well as again during inducer (MG132) treatment. Briefly, 24 hours after seeding the cells, the media was removed and replaced with media supplemented with 100 uM biotin followed by transfection. After 24 hours of transfection, the media was removed and replaced with biotin (100uM) as well as MG132 (10uM) supplemented complete media and incubated at 37°C, 5% CO_2_ incubator for times as indicated in figures.

### Biotin Labeling for TRAF6

For TRAF6 100 µM biotin was added for 15 min before harvest.

### PR-619 treatment

For treatment of PR-619, the cells were first treated with the inducer, in presence of 100uM biotin. After 2 hours, 10uM PR-619 was added to the media. Equal volume of DMSO was added in mock-treated cells. After 4 hours the cells were harvested.

For Ub-POD-TRAF6 experiments cells were treated for 15 min with 10 µM PR-619 in presence of 100 µM biotin.

### Immunoblotting

After required treatments, the harvested cells were lysed in 1X cell lysis buffer [50 mM Tris-HCl, pH 7.5, 150 mM NaCl, 1% Triton X-100, 0.1%SDS, 250u/ul benzonase, 1mM EGTA, 50mM NEM, protease inhibitor cocktail (Sigma Aldrich)] for 30 minutes at 4□°C. After pelleting the debris, the protein concentrations in the supernatant were measured using a BCA assay (ThermoFisher Scientific). The cell lysates were run on 4-20% Tris glycine gradient gels (Biorad) by SDS-PAGE followed by transfer to polyvinylidenedifluoride (PVDF) membranes. After the transfer, the membranes were incubated in blocking buffer (5% BSA in 1X PBS) for 30 min at room temperature followed by overnight incubation with the antibodies listed in table 2 at 4□°C. Next day, the membranes were incubated with fluorescent tagged secondary antibodies for 45 minutes at RT followed by imaging on a fluorescence scanner imaging system (Biorad).

### Streptavidin immunoprecipitation and immunoblotting

After required treatments, the harvested cells were lysed in 1X cell lysis buffer [50 mM Tris-HCl, pH 7.5, 150 mM NaCl, 1% Triton X-100, 0.1%SDS, 250u/ul benzonase, 1mM EGTA, 50mM NEM, protease inhibitor cocktail (Sigma Aldrich)] for 30 minutes at 4□°C. After pelleting the debris, the protein concentrations in the supernatant were measured using a BCA assay (ThermoFisher Scientific). For Streptavidin immunoprecipitation, the Streptavidin agarose beads (Thermo) were washed twice with the 1X cell lysis buffer and incubated with 1mg of lysed cell extracts for 2 h at 4°C subjected to end-to-end rotation. The beads were washed twice in wash buffer 1 (50 mM Tris-HCl, pH 7.5, 500 mM NaCl, 1%SDS, 1% Triton X-100, 10mM DTT, 1 mM EGTA, 50 mM NEM) and twice more with wash buffer 2 (50 mM Tris-HCl, pH 7.5, 150 mM NaCl, 0.1%SDS, 1% Triton X-100, 10mM DTT, 1 mM EGTA, 50 mM NEM). The final wash was done in wash buffer 3 (50 mM Tris-HCl, pH 7.5, 150 mM NaCl, 50mM NEM). All the washing steps were done for 5 min on a rotation wheel. Proteins were eluted by boiling with 2X gel loading dye. 10% of the elute was kept for immunoblotting and rest was kept for proteomics analysis. Input lysates were also boiled with 2X gel loading dye. 20ug of input cell lysates and 10ul of immunoprecipitated elutes were run on 4-20% Tris glycine gradient gels (Biorad). For detecting protein biotinylation, the proteins were transferred on a low fluorescence PVDF membrane followed by incubation with fluorescent tagged Streptavidin antibody (1:20,000 dilution in 1X PBS containing 5% BSA, 0.1% Tween-20, 0.1%SDS) for 45 min at RT. The membranes were washed three times, each for 5 min, with 1X PBS containing 0.2% Tween-20, and then once more with 1X PBS. The membranes were scanned on a fluorescence scanner imaging system (Biorad).

### Immunostaining, confocal microscopy and statistical analysis

HEK-293 cells were seeded on 22 mm coverslips coated with poly D lysine in a 6 well plate. The cells were co-transfected with HA-BirA/HA-BirA-RAD18 and (-2)AP-Ub constructs in presence of biotin (100uM). After 16 hours, the cells were exposed to 10 mJ/cm^2^ of UV and further incubated for 6 hours in biotin (100uM) containing complete media. The cells were then washed 5 times with 1X PBS and fixed with 4% PFA for 20 minutes at RT. After washing 3 times with 1X PBS, cells were blocked and permeabilized in solution 1 (5% BSA, 0.3% Triton X 100 in 1X PBS) for 3 hours at RT. After washing once with 1X PBS, the cells were incubated overnight with anti HA tag primary antibody (1:800) (Supplementary figure 4), anti-phospho Histone H2AX (Ser139) (1:400) (Figure 2E) in solution 2 (3% BSA, 0.1% Triton X 100 and 1X PBS). Next day cells were washed 3 times with solution 2 and 2 times with 1X PBS and further incubated with Streptavidin conjugated to Irdye 680 (1:200) and Alexa488 conjugated anti-rabbit Secondary antibody (1:200) for 1 hour in the dark in a humidified 37°C incubator. The cells were then washed 5 times with solution 2 and once with 1X PBS before mounting with ProLong antifade mountant with DAPI. Imaging was done in a Leica TCS SP5 confocal microscope (63X oil-immersion objective) keeping all the parameters same throughout the imaging process.

For statistical analysis, total no. of transfected cells and the no. of transfected cells with UV-irradiation induced RAD18 foci were counted from 6 different fields for each condition. Percentage difference of transfected cells with UV-irradiation induced RAD18 foci between HA-BirA-RAD18 and HA-BirA counterparts were calculated using Graphpad prism 9. Statistical significance of data was calculated by an unpaired T test and is marked as **** for p<0.0001.

To determine colocalization between Streptavidin (red) and HA-tag (green) images were analysed in FIJI (coloc tool). Results are indicative of 20 cells counted from 10 different images; error bars indicate standard deviation (p<0.0001, calculated by unpaired T test).

Total no. of HA-BirA/HA-BirA-Rad18 transfected cells and the no. of transfected cells showing colocalization between Streptavidin and phospho histone H2AX (Ser139) was calculated from 10 different fields and represented as a percentage. Statistical significance of data was calculated by an unpaired T test and is marked as **** for p<0.0001.

### BioID

For RAD18 and CHIP, the transfection, biotin treatment and inducer treatment regimes were followed as Ub-POD. After required treatments cells were lysed in 1X cell lysis buffer (50 mM Tris pH 7.4, 1% TritonX100, 150 mM NaCl, 0.4% SDS, 5 mM EDTA, 1 mM DTT, 1x complete protease inhibitor). The cell lysates were passed for 10–20 times (five to ten strokes) through a 25 G needle. Followed by sonication in a cold (4 °C) water bath for 5 minutes. The lysates were centrifuged at 16,000 x g, 10 min, 4°C. Protein concentration was measured by BCA assay and 2 mg protein was used for the streptavidin pulldown. For streptavidin pulldown, the streptavidin beads were washed by gently mixing with equilibration buffer (50 mM Tris pH 7–4, 150 mM NaCl, 1% Triton X-100, and 1 mM DTT). The 2mg lysate was incubated with the streptavidin beads for 2 hours at 4 °C on a rotating wheel. The beads were washed for 8 minutes on a rotation wheel with the following wash buffers: twice with wash buffer 1 (2% SDS in water), once with wash buffer 2 (50 mM HEPES pH 7.4, 1 mM EDTA, 500 mM NaCl, 1% Triton X-100, and 0.1% Na-deoxycholate). once with wash buffer 3 (10 mM Tris pH 8, 250 mM LiCl, 1 mM EDTA, 0.5% NP-40, and 0.5% Na-deoxycholate), twice with wash buffer 4 (50 mM Tris pH 7.4, 50 mM NaCl, 0.1% NP-40). 2X SDS sample buffer was added to the beads for elution followed by incubation at 95 °C for 15 minutes. The eluted samples were used for further analysis.

### Mass spectrometry and data analysis

Reduction of disulphide bridges in cysteine containing proteins was performed with dithiothreitol (56°C, 30 min, 10 mM in 50 mM HEPES, pH 8.5). Reduced cysteines were alkylated with 2-chloroacetamide (room temperature, in the dark, 30 min, 20 mM in 50 mM HEPES, pH 8.5). Samples were prepared using the SP3 protocol(*78*, *79*) and trypsin (sequencing grade, Promega) was added in an enzyme to protein ratio 1:50 for overnight digestion at 37°C. Next day, peptide recovery was done in HEPES buffer by collecting supernatant on magnet and combining with second elution wash of beads with HEPES buffer. Peptides were labelled with TMT10plex(*80*) Isobaric Label Reagent (Thermo Fisher) according to the manufacturer’s instructions. Samples were combined for the TMT10plex and for further sample clean up an OASIS® HLB µElution Plate (Waters) was used.

Offline high pH reverse phase fractionation was carried out on an Agilent 1200 Infinity high-performance liquid chromatography system, equipped with a Gemini C18 column (3 μm, 110 Å, 100 x 1.0 mm, Phenomenex)(*81*).

An UltiMate 3000 RSLC nano LC system (Dionex) fitted with a trapping cartridge (µ-Precolumn C18 PepMap 100, 5µm, 300 µm i.d. x 5 mm, 100 Å) and an analytical column (nanoEase™ M/Z HSS T3 column 75 µm x 250 mm C18, 1.8 µm, 100 Å, Waters) was used. Trapping was carried out with a constant flow of trapping solution (0.05% trifluoroacetic acid in water) at 30 µL/min onto the trapping column for 6 minutes. Subsequently, peptides were eluted via the analytical column running solvent A (0.1% formic acid in water, 3% DMSO) with a constant flow of 0.3 µL/min, with increasing percentage of solvent B (0.1% formic acid in acetonitrile, 3% DMSO) from 2% to 8% in 6 min, then 8% to 28% for a further 66 min, in another 4 min. from 28% to 38%, followed by an increase of B from 38-80% for 3min. and a re-equilibration back to 2% B for 5min. The outlet of the analytical column was coupled directly to an Orbitrap Fusion™ Lumos™ Tribrid™ Mass Spectrometer (Thermo) using the Nanospray Flex™ ion source in positive ion mode. The peptides were introduced into the Orbitrap Fusion Lumos via a Pico-Tip Emitter 360 µm OD x 20 µm ID; 10 µm tip (New Objective or CoAnn Technologies) and an applied spray voltage of 2.4 kV. The capillary temperature was set at 275°C. Full mass scan was acquired with mass range 375-1500 m/z in profile mode in the orbitrap with resolution of 120000. The filling time was set at maximum of 50 ms with a limitation of 4×105 ions. Data dependent acquisition (DDA) was performed with the resolution of the Orbitrap set to 30000, with a fill time of 94 ms and a limitation of 1×105 ions. A normalized collision energy of 38 was applied. MS2 data was acquired in profile mode.

IsobarQuant(*82*), Mascot (v2.2.07) and MS Fragger were used to process the acquired data. For MS Fragger, all raw files were converted to mzmL format using MSConvert from Proteowizard59, using peak picking from the vendor algorithm. For IsobarQuant and Mascot, the acquired data was searched against Uniprot Homo sapiens proteome (UP000005640) containing common contaminants and reversed sequences. For MS Fragger, files were searched using MSFragger v3.7 (PMID: 28394336) in Fragpipe v19.1 against the Swissprot Homo sapiens (UP000005640) database (20,594 entries) containing common contaminants and reversed sequences. The following modifications were included into the search parameters:

Carbamidomethyl (C) and TMT10 (K) (fixed modification), Acetyl (Protein N-term), Oxidation (M) and TMT10 (N-term) (variable modifications). For the full scan (MS1) a mass error tolerance of 10 ppm and for MS/MS (MS2) spectra of 0.02 Da was set. Further parameters were set: Trypsin as protease with an allowance of maximum two missed cleavages: a minimum peptide length of seven amino acids; at least two unique peptides were required for a protein identification. The false discovery rate on peptide and protein level was set to 0.01. GO-BP/CC/MF annotation analysis as well a GO-pathway enrichment analysis of the identified hits were performed in SRPlot (*83*).

### Statistical analysis

All experiments were independently reproduced at least two times with similar results obtained. No data were excluded from analysis. Statistical analysis for confocal images was performed by blinding the investigators to group allocation. Data are presented as the mean□±□s.d. for bar graphs. P□values were calculated using unpaired Student’s t-test using graphpad prism v.9.

## Methods for the TRAF6 Ub-POD

### Immunoblotting

For total lysates, cells were lysed in RIPA-buffer (50mM Tris, 150mM NaCl, 0.1% SDS, 0.5% sodium deoxycholate, 1% Triton-X, 50 µM PR-619, 1× protease inhibitors (Roche) and 1× PhosStop (Roche)) for 30 min on ice. After clearance by centrifugation at 20,000 × g for 10 min at 4 °C, protein concentrations were adjusted using BCA assay. Afterwards, lysates were boiled in sample buffer (200mM Tris-HCL, 6% SDS, 20% Glycerol, 0.1 g/ml DTT and 0.01mg Bromophenol Blue) for 5 min at 95 °C. Proteins were separated by SDS-PAGE and transferred onto nitrocellulose membranes. Membranes were blocked in 5% milk or 3% BSA in TBS supplemented with 0.1% Tween-20 (TBS-T) respectively. Membranes were incubated with primary antibodies overnight at 4 °C in blocking buffer, washed three times with TBS-T, incubated with secondary HRP-conjugated antibodies (1:10,000) for 1 h at room temperature, washed again with TBS-T and immediately analyzed by enhanced chemiluminescence.

### HA-IP

Briefly, cells expressing HA-constructs were washed 2 times with ice-cold DPBS, scraped in PBS and pellets resuspended and incubated in MCLB buffer (20 mM Tris-HCl pH 7.4, 150 mM NaCl, 0.5% NP-40, 1× protease inhibitors (Roche)) for 30 min on ice. After clearance by centrifugation at 20,000 × g for 10 min at 4 °C, protein concentrations were adjusted using BCA assay. Input samples were mixed with sample buffer and cooked for 5 min at 95 °C. IP was performed with preequilibrated HA agarose (Pierce) on an overhead rotator overnight at 4 °C. Next day, beads were washed 5 times with MCLB buffer. Iped proteins were eluted from beads by boiling in sample buffer for 5 min at 95 °C.

### Streptavidin pulldown and sample processing for mass spectrometry

After respective treatment, cells were washed 2 times with ice-cold DPBS, scraped in PBS and pellets either processed immediately or stored at -80°C. Cells were lysed in RIPA buffer for 30 min on ice. After clearance by centrifugation at 20,000 × g for 10 min at 4 °C, protein concentrations were adjusted using BCA assay. Cleared and adjusted supernatants were incubated overnight on an overhead rotator with preequilibrated streptavidin agarose (Sigma). Next day, beads were washed 2 times with RIPA buffer, 4 times with freshly prepared 3 M Urea buffer (Urea in 50 mM ammonium bicarbonate) and suspended in a defined volume of 3 M Urea buffer. Samples were reduced with 5 mM TCEP (Sigma) at 55 °C for 30 min, alkylated with 10 mM IAA (Sigma) at room temperature for 20 min and quenched with 20 mM DTT (Sigma). Samples were washed two times with freshly prepared 2 M Urea buffer (Urea in 50 mM ammonium bicarbonate), suspended in 50 μl of 2 M Urea buffer and digested with 1 μg trypsin per sample at 37 °C overnight. Peptides were collected by pooling the supernatant with two 50 μl 2 M Urea buffer washes, immediately acidified with 1% trifluoroacetic acid and concentrated by vacuum centrifugation. Digested peptides were desalted on custom-made C18 stage tips and reconstituted in 0.1% formic acid.

### MS data collection and analysis

Peptides were separated using an Easy-nLC1200 liquid chromatograph (Thermo Scientific) followed by peptide detection on a Q Exactive HF mass spectrometer (Thermo Scientific). Samples were separated on a 75 μm× 15 cm custom-made fused silica capillary packed with C18AQ resin (Reprosil-PUR 120, 1.9 μm, Dr. Maisch) with a 35 min acetonitrile gradient in 0.1% formic acid at a flow rate of 400 nl/min (5–38% ACN gradient for 23 min, 38–60% ACN gradient for 3 min, 60–95% ACN gradient for 2 min). Peptides were ionized using a Nanospray Flex Ion Source (Thermo Scientific). Peptides were identified in fullMS / ddMS² (Top15) mode, dynamic exclusion was enabled for 20 s and identifications with an unassigned charge or charges of one or >8 were rejected. MS1 resolution was set to 60,000 with a scan range of 300–1650 m/z, MS2 resolution to 15,000. Data collection was controlled by Tune/Xcalibur (Thermo Scientific). Raw data were analyzed using MaxQuant’s (version 1.6.0.1) Andromeda search engine in reversed decoy mode based on a human reference proteome (Uniprot-FASTA, UP000005640, downloaded June 2023) with an FDR of 0.01 at both peptide and protein levels. Digestion parameters were set to specific digestion with trypsin with a maximum number of 2 missed cleavage sites and a minimum peptide length of 7. Oxidation of methionine and amino-terminal acetylation were set as variable and carbamidomethylation of cysteine as fixed modifications. The tolerance window was set to 20 ppm (first search) and to 4.5 ppm (main search). Label-free quantification was set to a minimum ratio count of 2, re-quantification and match-between runs was selected and at least 4 biological replicates per condition were analyzed. Resulting protein group files were processed using Perseus (version 1.6.14.0). Briefly, common contaminants, reverse and site-specific identifications as well as proteins identified with a peptide count and MS/MS count <2 were excluded. Protein groups considered for statistical testing had further to be identified in at least 3 out of 4 replicates in at least one condition. After filtering, missing values were imputed from a normal distribution with a width of 0.3 relative to the standard deviation of measured values and a downshift of 1.8 standard deviation units of the valid data. Subsequent, two-sided t-test with multiple comparison correction (Permutation based-FDR) was performed. DAVID (version 2021) was used for functional enrichment analysis.

